# HyPR-seq: Single-cell quantification of chosen RNAs via hybridization and sequencing of DNA probes

**DOI:** 10.1101/2020.06.01.128314

**Authors:** Jamie L. Marshall, Benjamin R. Doughty, Vidya Subramanian, Qingbo Wang, Linlin M. Chen, Samuel G. Rodriques, Kaite Zhang, Philine Guckelberger, Charles P. Fulco, Joseph Nasser, Elizabeth J. Grinkevich, Teia Noel, Sarah Mangiameli, Anna Greka, Eric S. Lander, Fei Chen, Jesse M. Engreitz

## Abstract

Single-cell quantification of RNAs is important for understanding cellular heterogeneity and gene regulation, yet current approaches suffer from low sensitivity for individual transcripts, limiting their utility for many applications. Here we present Hybridization of Probes to RNA for sequencing (HyPR-seq), a method to sensitively quantify the expression of up to 100 chosen genes in single cells. HyPR-seq involves hybridizing DNA probes to RNA, distributing cells into nanoliter droplets, amplifying the probes with PCR, and sequencing the amplicons to quantify the expression of chosen genes. HyPR-seq achieves high sensitivity for individual transcripts, detects nonpolyadenylated and low-abundance transcripts, and can profile more than 100,000 single cells. We demonstrate how HyPR-seq can profile the effects of CRISPR perturbations in pooled screens, detect time-resolved changes in gene expression via measurements of gene introns, and detect rare transcripts and quantify cell type frequencies in tissue using low-abundance marker genes. By directing sequencing power to genes of interest and sensitively quantifying individual transcripts, HyPR-seq reduces costs by up to 100-fold compared to whole-transcriptome scRNA-seq, making HyPR-seq a powerful method for targeted RNA profiling in single cells.

## INTRODUCTION

Single-cell RNA sequencing (scRNA-seq) has emerged as a powerful and flexible tool for characterizing biological systems^1–5^. The ability to measure gene expression in thousands of single cells at once has enabled identifying and characterizing cellular heterogeneity in tissues, defining gene programs in previously uncharacterized cell types, and studying the dynamics of gene expression^6^. scRNA-seq has also facilitated high-throughput genetic screens that link CRISPR perturbations to effects on transcriptional programs in single cells^7–9^.

However, while scRNA-seq addresses certain biological questions by capturing an unbiased representation of the transcriptome, the utility of this approach is limited for applications that require sensitive detection of specific transcripts of interest. Such applications include profiling the dynamics of low-abundance or non-polyadenylated transcripts, quantifying rare cell types or states by measuring pre-defined marker genes, and characterizing the effects of perturbations to *cis*-regulatory elements on nearby genes. Addressing these questions requires considering the efficiency of detection for individual RNA transcripts (which can range from 5 to 45% for whole-transcriptome scRNA-seq sequenced to saturation^10–12^) and the depth of sequencing required to sample genes of interest (which might require hundreds of thousands of reads per cell for lowly expressed transcripts). To address the latter, previous studies have introduced several strategies to enrich and quantify specific transcripts in single cells, for example using target-specific reverse transcription of RNA during library preparation^13–15^ or hybrid selection, PCR, or linear amplification on single-cell cDNA libraries^16–19^. However, each of these approaches has certain limitations, such as the number of transcripts that can be measured, the inability to detect nonpolyadenylated transcripts, or the feasibility of profiling very large numbers of cells (see **Note S1**). New approaches are needed to quantify specific genes of interest in tens of thousands of single cells in a cost-effective and sensitive manner.

We developed a novel approach called Hybridization of Probes to RNA sequencing (HyPR-seq) that enables targeted quantification of RNAs in thousands of single cells. Our approach builds on the observation that single molecule fluorescence *in situ* hybridization (smFISH) — involving hybridization of labeled probes to target RNAs — can detect both polyadenylated and nonpolyadenylated RNAs with very high sensitivity^20–24^. We sought to develop a method that quantifies the same hybridization probes via next-generation DNA sequencing, rather than via imaging, to enable simultaneous readouts of hundreds of probes in thousands of single cells.

To do so, HyPR-seq involves hybridizing single-stranded DNA (ssDNA) probes to one or more RNA transcripts of interest, encapsulating single cells in 1-nanoliter droplets, PCR-amplifying the ssDNA probes, and sequencing these amplicons to quantify gene abundance in each cell. This method has greater than 20% sensitivity for individual transcripts, can simultaneously measure up to 100 polyadenylated and/or non-polyadenylated transcripts, and can profile more than 100,000 cells in a cost-effective manner. We demonstrate the utility of this approach by measuring the *cis*-regulatory effects of CRISPR perturbations to noncoding DNA elements, detecting nonpolyadenylated intronic RNA to measure time-resolved changes in gene expression, and quantifying the proportions of cell types in kidney tissue using low-abundance marker genes. By increasing the sensitivity of RNA detection while decreasing both sequencing and reagent costs, HyPR-seq extends the toolkit of single-cell methods to enable new experiments to investigate gene regulation and cellular programs.

## RESULTS

We began by adapting probes from the hybridization chain reaction (HCR) smFISH protocol^24,25^ for a sequencing-based readout. In HCR, two “initiator” probes anneal adjacent to each other on a target RNA molecule and are recognized by a metastable “hairpin” oligo, triggering a chain reaction in which fluorescently labeled oligos bind and extend (**Fig. S1**). In HyPR-seq, we eliminated the chain reaction and instead hybridize and ligate a single “readout” oligo to one of the initiator probes (**Fig. 1A**, **Fig. S1**). (We retain the cooperative initiator binding and metastable hairpin to increase the specificity of hybridization.) This ligation creates a ssDNA fragment that can be amplified by PCR and quantified by high-throughput sequencing (**Fig. S2**). We include a unique molecular identifier (UMI) on the amplified initiator probe to identify sequencing reads that originate from a single hybridization and ligation event.

**Figure 1.**
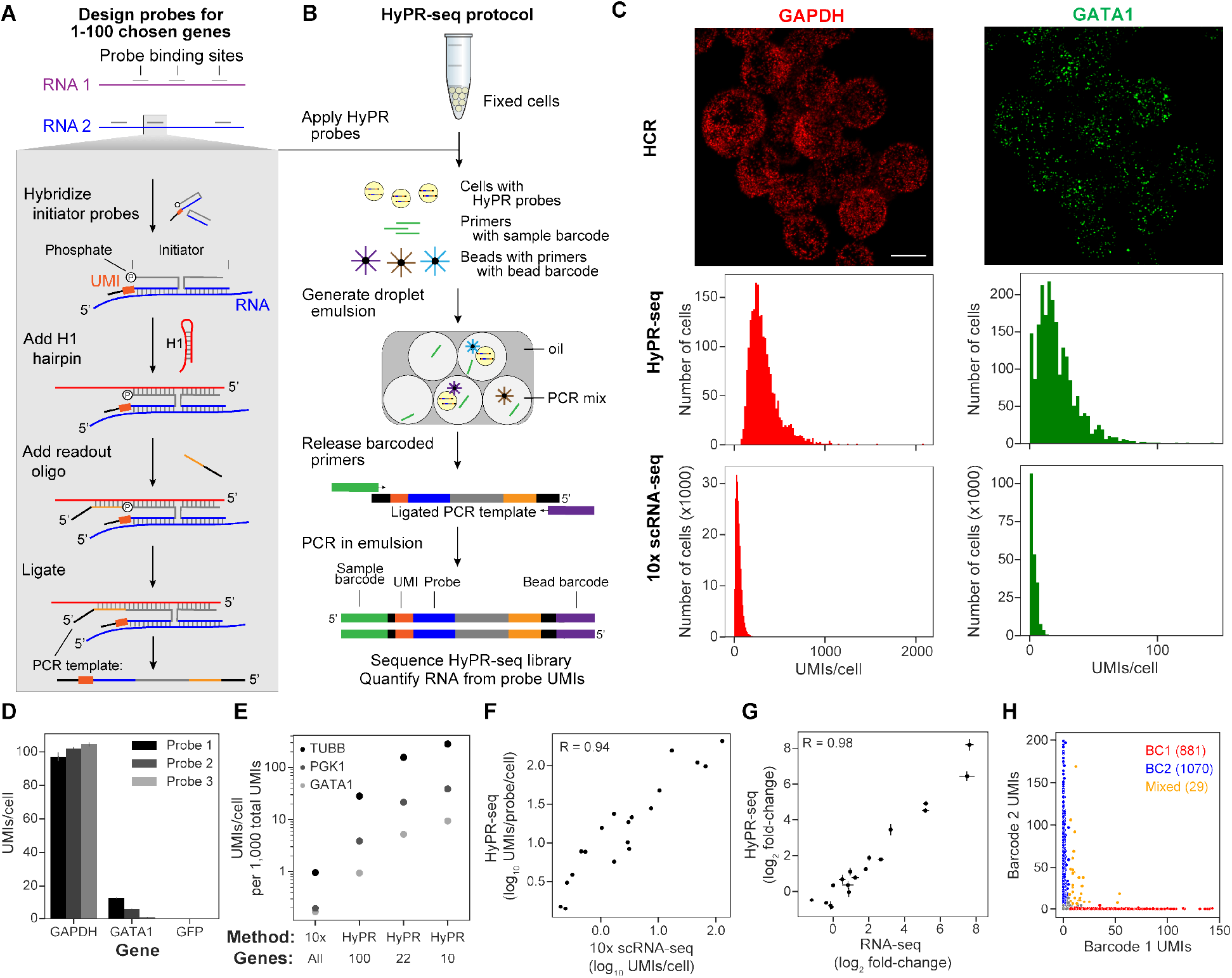
HyPR-seq enables single-cell quantification of selected RNAs. **A.** At a probe binding site, initiator probes bind adjacent to one another on an RNA of interest, creating a binding site for a hairpin oligo. A final readout oligo is annealed and ligated, forming a template for PCR amplification containing transcript information, unique molecular identifier (UMI), and handles for PCR and sequencing. **B.** Experimental workflow for HyPR-seq. **C.** Comparison of three single-cell methods in K562 cells. (Top) HCR measurements of GAPDH and GATA1 using probes targeting the same binding sites used for HyPR-seq. (Middle) Histogram of UMIs per cell for three probes targeting GAPDH and GATA1 from HyPR-seq and (bottom) 10x Genomics Chromium 3’ scRNA-seq. Scale bar represents 10μm, all experiments performed in K562 cells. **D.** HyPR-seq counts per cell for each of the 3 probes targeting GAPDH (left), GATA1 (center), or GFP (not expressed, right). Error bars represent 95% CI of the mean from two replicates. **E.** Counts per cell at a constant depth of 1,000 total UMIs per cell for 3 genes (GATA1, PGK1, and TUBB) as assayed by 10x Genomics Chromium 3’ scRNA-seq^26^ and HyPR-seq. Counts for HyPR-seq are shown for the 22 gene experiment in K562 cells and 2 simulated experiments (based on these counts) measuring different numbers of total genes. **F.** Counts per cell for 19 genes in K562 cells as measured by 10x Genomics Chromium 3’ scRNA-seq (X axis) and HyPR-seq (Y axis). Counts were normalized to the average probe for each gene, as genes were targeted by different numbers of probes **G.** In THP1 cells +/− LPS, fold changes of 18 genes assayed by bulk RNA-seq (X axis) and HyPR-seq (Y axis). Error bars represent 95% CI of the mean from three (RNA-seq) or four (HyPR-seq) replicates. **H.** To assess mixing, K562 cells were transduced to express two different barcode RNAs, each detectable by three HyPR probes. Scatterplot shows UMI counts for each barcode per droplet. Unassigned cells are in gray.

To use these HyPR-seq probes to detect RNAs in single cells (**Fig. 1B**), we: (i) crosslink and permeabilize a population of cells, (ii) hybridize initiator probes to 1-100 target RNAs, and (iii) hybridize and ligate the hairpin and readout oligos. We then (iv) distribute single cells and DNA-barcoded microparticles (“beads”) into an emulsion PCR using a commercially available automated microfluidic droplet-maker. Each bead carries photocleavable primers with a unique clonal barcode. We then (v) cleave primers off of the beads using ultraviolet light, (vi) PCR-amplify the HyPR-seq probes in emulsion, (vii) sequence the resulting amplicons, and (viii) quantify the expression of each target gene based on the UMI counts of corresponding initiator probes. During the microfluidic emulsion step, we Poisson-load cells and beads into droplets, expecting 5-15% of droplets to contain 1 cell and at least 1 bead. Droplets containing more than 1 bead are computationally identified and merged (see Methods, **Fig. S3**). In our implementation, each sample from the droplet-maker yields approximately 500-2,500 single cells. Our complete HyPR-seq probe design and data analysis pipeline is available at https://github.com/EngreitzLab/hypr-seq.

To test the performance of HyPR-seq, we designed and applied 102 HyPR-seq probes to detect 22 genes (2-10 probes per gene, **Table S1**) in K562 human leukemia cells in two biological replicates. We sequenced the libraries to saturation (17,111 reads per cell) and identified 4,605 total cells, with an average of 4,021 UMIs per cell (**Fig. S4A**, **Table S2**). UMI counts per probe per cell were highly reproducible between biological replicates (Pearson’s R > 0.99; **Fig. S4B**).

### Specificity

To investigate the specificity of HyPR-seq, we first compared probes targeting two highly expressed genes (GAPDH and GATA1) to control probes that did not target a gene expressed in K562 cells. The GAPDH and GATA1 probes gave 10,000-fold and 500-fold higher signal compared to the control probes: for GAPDH, we measured an average of 303.7 UMIs per cell (101.2 UMIs per probe per cell); for GATA1, an average of 19.1 UMIs per cell (6.4 UMIs per probe per cell); and for the control probes, an average of fewer than 0.03 UMIs per cell (<0.01 UMIs per probe per cell) (**Fig. 1C, D**). As expected, the 3 independent probes for each gene (targeting different locations on the same mRNA) varied in their UMI counts, presumably due to sequence-specific differences in hybridization efficiency (**Fig. 1D**). Across all genes in the experiment, ~75% of probes had UMI counts within 2-fold of the median probe for a given gene (**Fig. S4C**) and yielded counts >100-fold above the negative control probes. We estimate that HyPR-seq can specifically detect transcripts expressed above approximately 1 TPM (**Fig. S4D**, see Methods).

### Detection rate and sequencing efficiency

We evaluated the counts per gene observed in HyPR-seq compared to scRNA-seq and smFISH experiments in K562 cells (**Fig. 1C**). We first counted UMIs in this HyPR-seq experiment (sequenced to saturation, ~4,000 UMIs per cell) compared to a 10x Genomics Chromium 3’ scRNA-seq dataset^26^ (not sequenced to saturation, ~18,000 UMIs per cell). We observed an average of 36-fold more UMIs per gene in HyPR-seq than in scRNA-seq (**Fig. 1C**, **Fig. S4E**), corresponding to 162-fold more UMIs per cell per 1,000 total UMIs. This enrichment will vary between 30- and 300-fold depending on the number of genes targeted in the HyPR-seq experiment (**Fig. 1E**). We then compared HyPR-seq measurements (sequencing UMIs) to smFISH of GATA1 mRNA (counting spots), and found that the best HyPR-seq probe targeting GATA1 yielded 20% of the smFISH counts, and together three HyPR-seq probes targeting GATA1 yielded 31% of the smFISH counts (**Fig. S4F**). In smFISH, 20-50 probes per gene are used per RNA to obtain near quantitative detection efficiency (~90%), assuming that probes bind independently to their target RNAs^27^. Similarly, for HyPR-seq, increasing the number of probes per gene will increase detection efficiency for genes expressed above the specificity limit of 1 TPM. The ability to adjust the number of probes per gene will allow users to tune the detection rate and fraction of sequencing reads devoted to different genes.

### Quantification accuracy

We examined the accuracy of HyPR-seq in quantifying the expression levels of different genes in a single condition (“absolute quantification”) and quantifying the foldchange in expression of a given gene across conditions (“relative quantification”).

To assess absolute quantification, we compared our HyPR-seq data across 19 genes (not including GFP, BFP, and the non-coding RNA PVT1) to 10x Genomics Chromium 3’ scRNA-seq data collected for the same cell type^26^. UMI counts per probe per cell from HyPR-seq correlated well with scRNA-seq counts for the same genes (Pearson’s R = 0.94; **Fig. 1F**, see also **Fig. S4D**).

To assess relative quantification, we performed two experiments. First, we measured the expression of an RNA encoding blue fluorescent protein (BFP) under the control of a doxycycline-inducible promoter at 14 timepoints after induction, during which BFP mRNA increased >100-fold. HyPR-seq measurements of BFP mRNA expression correlated well with corresponding measurements by qPCR (Pearson’s R = 0.95; **Fig. S4G**). Second, we examined 18 genes whose expression levels are known to change between 0.5- and 250-fold in THP1 monocytic leukemia cells upon stimulation with bacterial lipopolysaccharide (LPS). The fold-change in gene expression in LPS-stimulated vs. unstimulated cells correlated well between HyPR-seq and bulk RNA-seq (Pearson’s R = 0.98; **Fig. 1G**), and these fold-changes were similar across most probes targeting the same gene (**Fig. S4H**). These results indicate that HyPR-seq accurately quantifies changes in gene expression across a wide dynamic range.

### Single-cell purity

We demonstrated the single-cell purity of HyPR-seq through a single-cell mixing experiment. We engineered two cell lines to each express a unique transcript detectable by HyPR-seq (“detection barcodes”, see Methods). We applied probes that recognized the two different detection barcodes to both cell lines, then mixed the cells before distributing them into emulsion droplets. We found that 98.5% of droplets contained UMIs from a single detection barcode and 1.5% of the droplets had UMIs from both barcodes. This suggests a cell doublet rate of approximately 2.9% (in a 50/50 mix of cells, half of the doublets will involve two cells with the same barcode), compared to a theoretical rate of about 0.9% based on the density of cell loading (**Fig. 1H**). These data indicate a high level of single-cell purity in the HyPR-seq emulsion PCR.

Having demonstrated the essential technical capabilities of HyPR-seq, we examined its utility in three areas where existing single-cell tools remain limited: (1) profiling the effects of CRISPR perturbations on individual genes of interest, (2) measuring non-polyadenylated RNAs such as gene introns, and (3) detecting rare cell types via lowly expressed marker genes in complex tissue.

### CRISPR screens with HyPR-Seq to map perturbation effects on gene expression

We explored the utility of HyPR-seq in generating low-cost readouts of CRISPR perturbations to DNA regulatory elements^26,28^, where sensitive gene detection is critical for obtaining quantitative readouts of the effects of these elements on nearby genes.

We first developed an approach to apply HyPR-seq to pooled CRISPR screens, which requires detecting which gRNAs are expressed in each single cell in a population. We designed a lentiviral vector that expresses a gRNA from a U6 promoter as well as a unique detection barcode from a Pol II promoter (**Fig. 2A**). We designed HyPR-seq probes to detect these barcodes and thereby determine which gRNA is expressed in a given cell. Each detection barcode includes three distinct probe binding sites, allowing us to combinatorially map a large number of unique gRNAs using a more limited set of validated probes.

**Figure 2.**
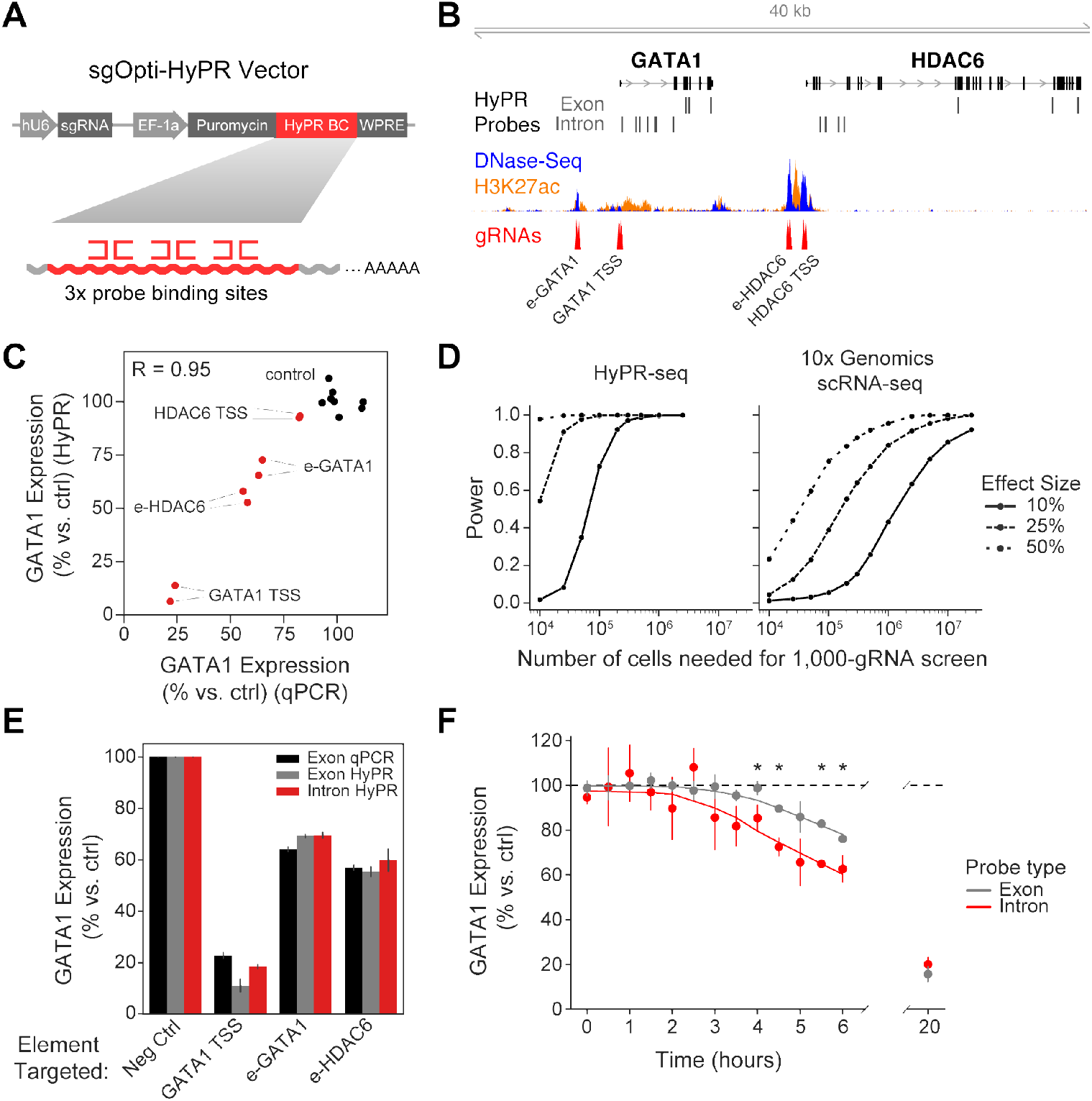
Application of HyPR-seq to CRISPR screens for mature mRNAs and introns. **A.** Design of sgOpti-HyPR barcoded vector. sgOpti vector was modified to express a detection barcode consisting of three probe binding sites (HyPR BC) from a Pol II promoter along with a sgRNA. **B.** Location of CRISPRi gRNAs and HyPR-seq probes in the *GATA1* locus, with DNase-seq and H3K27ac ChIP-seq signals from K562 cells. **C.** Relative GATA1 expression as measured by qPCR (X axis) and HyPR-seq (Y axis) for a small library of 16 gRNAs (8 non-targeting controls [black] and 2 targeting each of 4 elements in the *GATA1* locus [red]). qPCR measurements come from cell lines individually infected with each gRNA, while HyPR-seq measurements come from single cells in a pooled screen. **D.** Power to detect changes in gene expression of various magnitudes (Y axis, lines represent 10%, 25%, and 50% effect sizes) as a function of the total number of cells profiled (X axis). Power is shown for a hypothetical screen of 1,000 CRISPR perturbations and is averaged over 37 genes expressed at >1.5 UMIs per cell in a HyPR-seq dataset (left) and the same genes in a 10x Genomics Chromium 3’ scRNA-seq dataset (right) sequenced to ~18,000 UMIs per cell^26^. Power is estimated by testing the indicated number of cells (divided by 1,000) against M total cells (where M=5,000 for HyPR-seq and 200,000 for 10x Genomics Chromium 3’ scRNA-seq) to mimic a screen with an excess of non-targeting control gRNAs. **E.** GATA1 knockdown as measured by qPCR (with primers targeting the mature mRNA, black) or HyPR-seq with probes designed against the exons (gray) or first intron (red) of GATA1. Error bars represent 95% CI on the mean of two replicates. **F.** GATA1 expression in cells with gRNAs targeting *GATA1* TSS as measured by both exon- (gray) and introntargeting (red) probes over a 20-hour timecourse. Mean and 95% CI are shown for three replicates at each of 14 timepoints, and the curves are fit with lowess regression. Asterisks indicate a significant difference in the knockdown as measured by introns versus exons.

We tested this approach using a CRISPR interference (CRISPRi) screen in K562 cells to inhibit four elements in the *GATA1* locus that we previously found to regulate GATA1^29,30^ (**Fig. 2B**). We infected K562 cells expressing KRAB-dCas9 from a doxycycline inducible promoter with lentiviral constructs encoding 16 gRNAs along with linked detection barcodes (8 control gRNAs and 2 targeting each element). We robustly detected these barcodes (with at least 10 UMIs per cell in 90% of cells) and were able to assign a unique guide to >80% of cells (**Fig. S5A, B**). We quantified the effects of each gRNA on the expression of GATA1 and found that the changes in gene expression detected by HyPR-seq strongly correlated with those measured by qPCR (Pearson’s R = 0.96; **Fig. 2C**).

Such an approach could be useful for large-scale screens to measure the effects of enhancers on nearby genes, where existing approaches read out either effects on one gene at a time with very high sensitivity^30^ or effects on all genes in the transcriptome with low per-gene sensitivity^26,28^. Using data from our pilot experiments, we calculated the power of HyPR-seq to profile the effects of 1,000 CRISPR perturbations on 50 selected genes, the scale needed to systematically connect all putative enhancers in a genomic locus to their target genes^30^. For this experiment, HyPR-seq would require profiling approximately 25,000 cells at 5,000 reads per cell to achieve 90% power to detect 25% changes in expression for all genes expressed at >1 transcript per million (TPM, **Fig. 2D**, left). In contrast, whole-transcriptome scRNA-seq would require profiling >1,000,000 cells at 20,000 reads per cell to achieve similar power (**Fig. 2D**, right), increasing the total cost by two orders of magnitude. Thus, HyPR-seq could provide a powerful and cost-effective approach to profile the effects of CRISPR perturbations on a set of selected genes.

### Detecting non-polyadenylated introns to measure perturbation effects with increased temporal resolution

Due to its hybridization-based detection of RNA, HyPR-seq can, in theory, quantify non-polyadenylated transcripts that are difficult to detect with existing droplet-based scRNA-seq approaches. To demonstrate this, we used HyPR-seq to detect gene introns, which are present at lower copy numbers than mature mRNAs due to their short half-lives but whose abundance can be used to estimate gene transcription rates^22^. We designed probes targeting the introns of four genes (GATA1, HDAC6, MYC, and PVT1) and performed HyPR-seq in 4,605 K562 cells. We detected absolute signals that correlated with the transcription rates of these four genes as measured by PRO-seq (Pearson’s R = 0.99; **Fig. S5D**), including an average of 13.1 UMIs per cell for introns of MYC (a highly transcribed gene in K562 cells) and 3.5 UMIs per cell for introns of PVT1 (a less transcribed gene in K562 cells).

Quantifying intron abundance could enable more temporally precise measurements of the effects of perturbations on gene expression. We examined the effects of promoter or enhancer inhibition with CRISPRi on GATA1 expression over a 20-hour timecourse of KRAB-dCas9 induction in 112,056 K562 cells. We used exon-targeting probes to estimate steady-state mRNA levels and intron-targeting probes to estimate transcription rates. Both intron- and exon-targeted HyPR-seq probes showed equivalent levels of reduction after 20 hours, indicating that intron-targeting probes show similar quantitation and specificity (**Fig. 2E**). However, we detected a decrease in intron signal hours earlier than for exon signal, consistent with the shorter half-life of introns compared to mature mRNAs (**Fig. 2F**). Thus, HyPR-seq can detect time-resolved changes in gene expression via direct detection of low-abundance gene introns and may enable multiplexed detection of other non-polyadenylated RNAs that are difficult to capture using existing droplet-based single-cell methods.

### Measuring cell type frequencies and low-abundance genes in tissue

HyPR-seq could enable new types of highly multiplexed experiments to measure the expression of genes of interest in complex tissues. For example, determining the frequencies of closely related cell types in a tissue can require the detection of specific marker genes that may not be highly expressed, which is challenging with whole-transcriptome RNA-seq^31^.

To test this, we applied HyPR-seq to detect changes in cell type frequency in a mouse model of diabetic kidney disease (DKD)^32^. We designed HyPR-seq probes to distinguish 11 cell populations using 25 canonical marker genes. Some of these marker genes were lowly expressed in their corresponding cell types (<2 TPM) and are not robustly detected in existing scRNA-seq datasets (**Table S7**)^33–37^. Accordingly, we included up to 20 probe sets per gene to enable sensitive detection (see Methods). We applied HyPR-seq to dissociated single cells from the kidneys of 12-week old wild-type (BTBR *wt/wt*) and diabetic (BTBR *ob/ob*) mice and detected 29,125 single cells after doublet detection and filtering steps (see Methods). UMAP visualization of the HyPR-seq data identified 11 clusters, including all of the targeted cell populations (**Fig. 3A, B**, **Fig. S6**, and **Table S7**).

**Figure 3.**
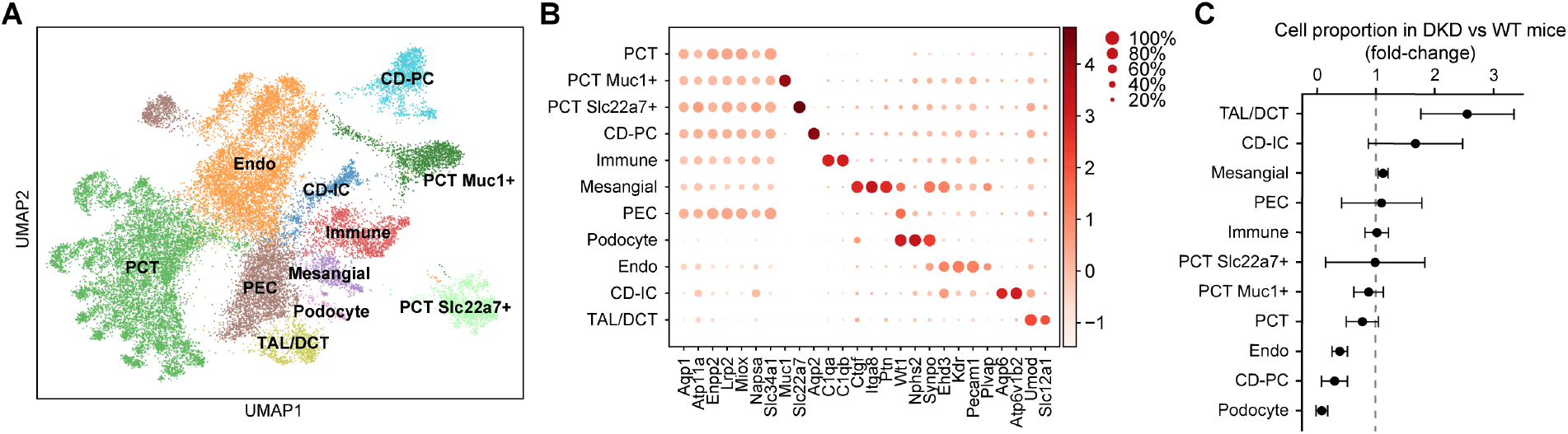
HyPR-seq in mouse kidney cells detects changes in cell frequencies in diabetes. **A.** UMAP visualization of HyPR-seq data for 31 genes in 14,288 BTBR *wt/wt* cells and 14,837 BTBR *ob/ob* cells identifies 11 distinct cell types. PCT: proximal-convoluted tubule. CD-PC: collecting duct principal cells. PEC: parietal epithelial cell. Endo: endothelial cell. TAL/DCT: thick ascending limb / distal convoluted tubule cell. **B.** Dot plot displays kidney cell types with the fraction of cells in each subtype expressing each gene (circle size) and the relative level of gene expression (color). **C.** Change in cell proportions in diabetic versus wild-type mice. Error bars represent the standard deviation of the ratio from two biological replicates (see Methods).

HyPR-seq accurately detected changes in cellular composition in DKD and enabled quantification of specific genes of interest with only an average of 203 UMIs per cell. For example, podocytes represent a rare cell type that plays a key role in the glomerular filtration barrier^38^. We observed podocytes – marked by the combined expression of *Wt1*, *Nphs2*, and *Synpo* – at 0.9% frequency in wild-type mice (132 out of 14,288 cells). Podocytes decreased in frequency by >10-fold in diabetic mice (to 12 out of 14,837 total cells, 0.08%), consistent with previous reports that podocytes are one of the earliest cell types to be damaged and lost in DKD (**Fig. 3C**)^39–41^. HyPR-seq also detected an increase in thick ascending limb/distal convoluted tubule (TAL/DCT) cells and a decrease in endothelial cells, in agreement with early changes observed in DKD (**Fig. 3C**)^42^. Finally, HyPR-seq detected various subtypes of epithelial cells, including two subclusters of proximal convoluted tubule cells (PCTs, Slc22a7+ and Muc1+) and collecting duct principal cells (CD-PCs, Aqp2+), and indicated that CD-PCs are reduced in frequency in DKD (**Fig. 3B, C**).

Together, these observations were possible using only 203 UMIs per cell, ~10-fold fewer than were used to make similar observations using scRNA-seq^42^. HyPR-seq will enable high-throughput experiments to measure cellular responses to environmental, chemical, or genetic perturbations in primary cells and complex tissues, including measuring individual genes of interest that may be lowly expressed such as drug targets, transcription factors, and signaling molecules.

## DISCUSSION

Here, we described HyPR-seq, a microfluidic droplet-based approach for cost-effective and sensitive profiling of a chosen subset of RNA molecules in single cells. HyPR-seq provides a unique combination of capabilities that overcome key limitations of existing single-cell techniques. First, by adapting *in situ* hybridization probes for a sequencing-based readout, HyPR-seq achieves a per-probe RNA detection sensitivity of approximately 20% relative to smFISH, while also expanding multiplexing (in this study, we included up to 250 probes in a single experiment). Second, HyPR-seq can easily be scaled to examine more than 100,000 cells in a single experiment, facilitating large screens. Third, HyPR-seq can detect RNA species that are not targeted by polyA-based scRNA-seq approaches, including introns, enabling time-resolved studies of transcription. Finally, HyPR-seq provides a cost-effective approach for targeted quantification of selected transcripts (up to 100) in single cells by reducing reagents costs per cell by 5-fold and sequencing costs per cell by up to 100-fold versus whole-transcriptome scRNA-seq.

HyPR-seq does have several acknowledged limitations. HyPR-seq does not sequence the RNA molecule itself, making it unsuitable for detecting RNA sequence variants or modifications. The current protocol involves multiple rounds of washes for probe hybridization and ligation, which results in some cell loss and requires starting an experiment with 1 million or more cells. Finally, because HyPR-seq involves hybridization of probes, it could be more difficult to detect certain RNA transcripts that have high sequence homology with others.

The unique capabilities of HyPR-seq will enable experiments that were previously impractical using existing tools. For example, HyPR-seq could allow for large-scale CRISPR-based studies to perturb hundreds of regulatory elements in a single locus and profile their effects on all nearby genes. We anticipate that HyPR-seq will be broadly useful for sensitively quantifying RNA expression across a wide range of systems to study gene regulation and cellular heterogeneity.

## MATERIALS AND METHODS

### Design of HyPR-seq probes

We adapted the hybridization chain reaction (HCR, version 3)^24^ probe design for our droplet-based HyPR-seq method (**Fig. S1, Fig. S2**).

#### Initiator probes

Two initiator probes, a 5’ probe and a 3’ probe, each contain 25bp of sequence homologous to the target RNA and other necessary sequences.

The structure of the 5’ probe is:

[5’ Initiator] [5’ Spacer] [5’ Homology] [UMI] [Primer Binding Site]

The structure of the 3’ probe is:

[3’ Homology] [3’ Spacer] [3’ Initiator]

The constant sequences are:

5’ Initiator: /5Phos/GGAGGGCAGCAAACGG

5’ Spacer: AA

UMI: NNNNNNNNNN

Primer Binding Site: CTCGACCGTTAGCAAAGCTC

3’ Spacer: TA

3’ Initiator: GAAGAGTCTTCCTTTACG

For probes in this study, we use the “B1” initiator system from HCR^24^. Compared to the HCR initiator probes, the modifications we made for HyPR-seq are entirely in the 5’ probe. We added a 5’ phosphate (for ligation in HyPR-seq), attached a 20-bp sequencing adapter (primer binding site) to the 3’ end, and removed 2 bp from the 5’ end of the initiator and added it to the readout oligo, which we found improved the specificity of HyPR-seq (data not shown).

Following hybridization of the initiator probes, we add the hairpin oligo B1H1 (CGTAAAGGAAGACTCTTCCCGTTTGCTGCCCTCCTCGCATTCTTTCTTGAGGAGGGCAGCAAACGGGAAGAG, Molecular Instruments).

Finally, we add a “readout” oligo adapted from HCR hairpin B1H2 (CTTACGGATGTTGCACCAGCAAGAAAGAATGCGA, IDT), which is ligated to the 5’ initiator probe (**Fig. S2**).

### Custom barcoded bead design

Custom 68-mer beads were ordered from Chemgenes on 10 micron Agilent polystyrene beads at an oligo synthesis scale of 10 μmole, using the following sequence:

5’ -bead-linker-PC-linker-

CAAGCAGAAGACGGCATACGAGATJJJJJJJJJJJJGTTGGCACCAGGCTTACGGATGTTG CACCAGC-3’.

### Selecting target sequences for HyPR-seq probes

HyPR-seq probes can target any site on an RNA that enables specific binding. We chose the 5’ and 3’ homology sequences on the initiator probes that, similar to HCR, target 25-bp regions on a transcript of interest separated by a 2-bp spacer (52 bp total). We developed a custom design script to identify 52 bp RNA homology sequences. We tiled candidate sequences across the RNA and excluded those that (i) contained homopolymer repeats (N>5), (ii) were predicted to form hairpins or dimers, (iii) had GC-content outside of 40-65%, (iv) contained more than 5 bases of repetitive sequence (by comparison to RepeatMasker^43^), or (v) had a match with >25% identity when compared with BLAST to the rest of the transcriptome (RefSeq). From the list of valid homology sequences, we generated the final list by selecting a smaller number of sequences (typically 4-6) spaced evenly across the transcript of interest. Then, the 52 bp homology sequence was split to produce the probe homology regions. The reverse complement of bases 1-25 (in the 5’-3’ direction on the RNA) form the 3’ homology sequence, and the reverse complement of bases 28-52 form the 5’ homology sequence, with bases 26-27 acting as an unbound spacer.

Probes targeting the introns of genes were designed similarly, except we confined our search for homology regions to the first 5kb of the first intron (or first 5 kb of any introns, if the first intron is shorter than 5 kb), in order to detect RNA species whose appearance would most closely correlate with the initiation of transcription.

### HyPR-seq Experimental Protocol

#### Cell preparation

For HyPR-seq, cells are harvested in 1X phosphate buffered saline (PBS) at 350g for 5 mins at 4°C in a swinging bucket rotor. Cells were fixed in 4% formaldehyde solution (4% formaldehyde in 1X PBS and 0.1% Tween 20) for one hour at room temperature with rocking at a concentration of 1 million cells per mL, with up to 10 million cells in a 15-mL conical tube. For all following washes and centrifugations after fixation, cells were spun at 850g for 5 mins at room temperature. Fixed cells were washed twice with 1X PBS containing 0.2% Tween 20 (1X PBST). The washed cells were then permeabilized in 70% ice cold ethanol at a concentration of 1M/mL, with up to 10 million cells in a 15-mL conical, and stored at 4°C for 10 mins. The permeabilized cells were harvested and washed twice with 1X PBST. Then, the cells were transferred to 2-mL round-bottom tubes for the remainder of the protocol, with up to 5 million cells per tube at a concentration of 10M/mL. The cells were resuspended in 500 μL of probe hybridization buffer (5X SSC, 30% formamide, 0.1% Tween 20) and incubated at 37°C for 5 mins. The pre-hybridized cells were centrifuged, resuspended in the probes mix (which was prepared by pooling probesets in probe hybridization buffer to a final concentration of 20 nM per probe), and incubated overnight at 37°C in a hybridization oven (VWR Cat# 10055-006). After hybridization, cells were harvested and washed in probe hybridization buffer for 10 minutes at 37°C. The wash step was repeated 3 additional times, for a total of 4 washes. In the meantime, snapcooled hairpin solutions were prepared as follows, for 5 million cells. In separate tubes, 5 μL each of the B1H1 hairpin and readout oligo (each at a concentration of 3 μM) were incubated at 95°C for 90 seconds, then allowed to cool to room temperature over 30 minutes. After the final wash in probe hybridization buffer, cells were resuspended in 5X SSCT (5X SSC, 0.1% Tween 20) and incubated at room temperature for 5 mins. Cells were centrifuged then resuspended in 75 nM snapcooled B1H1 hairpin in 5X SSCT (for 5 million cells, this was 200 μL 5X SSCT and 5 μL of the snapcooled B1H1 hairpin). The cells were incubated at 37°C for one hour in the hybridization oven. After the B1H1 hairpin hybridization, the cells were washed twice with 5X SSCT, then resuspended in 75 nM snapcooled readout oligo in 5X SSCT (for 5 million cells, this was 200 μL of 5X SSCT and 5 μL of the snapcooled readout oligo). Cells were incubated 37°C for one hour. After oligo incubation, cells were harvested and washed 3 times in 5X SSCT. After the last spin, cells were washed in 1X T4 Ligase reaction buffer (NEB #B0202S) before being resuspended in 1X ligase (1X T4 ligase buffer, 1:100 dilution of T4 DNA ligase NEB #M0202S) and incubated at room temperature for one hour. Following ligation, cells were washed three times in 1X PBST and filtered through a 20 μm filter (20μm pluriStrainer cat. 43-0020-01). The cells were then counted using a hemocytometer (Fisher Scientific SKU #DHCF015) and checked for single cell suspension before proceeding to droplet generation. For long-term storage of the cells, RNase inhibitor was added to the cell suspension (NEB #M0314S).

#### Generation of emulsions

To generate emulsions, we first combined 1X EvaGreen Supermix (Bio-Rad Cat #186-4033), 500 nM indexing primer (**Table S3**), 2,000 cells and 20,000 barcoded beads (Chemgenes, as described above) in a 20 μl reaction. Once the PCR mix is made, emulsions were generated using QX200 Digital Droplet Generator (Bio-Rad #1864002) as per manufacturer’s instructions. Briefly, the droplet generation cartridge (Bio-Rad #186-4007) was inserted into the holder (Bio-Rad #186-3051) and 20 μl of the prepared PCR mix was added to the middle sample well, as per manufacturer’s instructions. 70 μl of the droplet generating oil (Bio-Rad, #186-4005) was added into the bottom oil well. This was repeated for all wells in a chip, where unused sample wells were filled with 1X PBS. Then, the gasket (Bio-Rad #186-4007) was placed over the filled cartridge and the cartridge was placed in the droplet generator. Once the droplets were formed, the cartridge containing the emulsions was placed under the UV lamp (6.5 J/cm2 at 365 nm) about 3-8 cms away from the bulb for 5 mins. After UV exposure, ~50 μl of droplets per well was transferred to 96 well plates (Eppendorf #951020362) and sealed with foil (Bio-Rad #181-4040) using a plate sealer (PX1 Plate Sealer #181-4000) for droplet PCR amplification (Eppendorf Mastercycler Pro #E90030010). The following cycling conditions were used: Denaturation: 94°C for 30 sec; Cycling: 30 cycles of 94°C for 5 sec, 64°C for 30 sec, and 72°C for 30 sec; Final extension: 72°C for 5 min. All PCR cycling steps were performed with 50% ramp rate (2°C/min). To break and clean the emulsions, 1-4 PCR wells were combined in a tube and 40 μL of 97% 1H,1H,2H,2H-Perfluoro-1-octanol (Sigma-Aldrich #370533) was added. The tubes were vortexed for 5 sec to ensure complete emulsion breakage and spun at 1000g for 1 min. The top aqueous layer was carefully removed and cleaned with 1.8X SPRI beads according to manufacturer’s instructions. The amplicon libraries were loaded on a gel to determine size (206bp) and to ensure no primer dimers remained. The libraries were quantified by Qubit before proceeding to sequencing.

#### Sequencing

The libraries were loaded at a concentration of 6pM on a MiSeq and at 1.8pM on a NextSeq 550. The sequencing specifications were as follows: Read 1: 35bp, Index 1: 8bp and Index 2: 12bp (see also **Fig. S2**). Sequencing the HyPR-seq libraries on the MiSeq and NextSeq both required custom Read 1 (GACACATGGGCGGAGCTTTGCTAACGGTCGAG, IDT) and custom Index 1 primers (GCTGGTGCAACATCCGTAAGCCTGGTGCCAAC, IDT). The NextSeq additionally required a custom Index 2 primer to read the sample indices. (CTCGACCGTTAGCAAAGCTCCGCCCATGTGTC, IDT). All custom primers were added according to manufacturer’s instructions. The depth of sequencing scaled with the number of genes. For the K562 experiments described (targeting 22 highly expressed genes), we typically aimed to sequence 10,000 reads per cell x 1,000 cells per well = 10 million reads.

### HyPR-seq computational pipeline

We built a custom pipeline to analyze HyPR-seq data by taking raw sequencing reads and constructing a count matrix (counts per probe per cell). Briefly, we filter low-quality reads, identify real bead barcodes, map to our set of probes, remove PCR duplicates using UMIs, and combine data that came from multiple bead barcodes in the same droplet (see below). We have made our analysis pipeline available on GitHub (https://github.com/EngreitzLab/hypr-seq) and detailed the key steps below (standard settings used except where indicated):

1. We demultiplex reads by well barcode (index 2) into separate FASTQs per experiment.
2. We then filter reads, removing short reads in index 1 (bead barcode) and reads with low quality: fastp --length_required 12.
3. We build a whitelist of bead barcodes to include in downstream analyses. We turn off error correction for sequencing/PCR errors for adjacent bead barcodes and typically rely on the built-in method for finding the UMI threshold to distinguish real cells from background (“knee”): umi_tools whitelist --bc-pattern=NNNNNNNNNN --bc-pattern2=CCCCCCCCCCCC --method=umis --error-correct-threshold 0 --knee-method=distance. In some cases, we found that we needed to manually specify the knee threshold with -set-cell-number=[BB_NUM].
4. We extract bead barcodes and UMIs from the read: umi_tools extract --bc-pattern=NNNNNNNNNN --bc-pattern2=CCCCCCCCCCCC --filter-cell-barcode -- whitelist=[WHITELIST].
5. We trim the remaining sequence to 25bp matching the probe variable sequence: fastx_trimmer -f 1 -l 25 -z -Q33.
6. We map reads to a custom Bowtie index (generated with bowtie-build from a custom FASTA file with one contig per probe): bowtie -v 1.
7. We get sorted and indexed BAM files with SAMTools: view, sort, and index.
8. We construct a table of reads grouped by UMI, transcript, and bead barcode: umi_tools group --per-cell.
9. We feed the resulting (UMI, transcript, bead barcode) tuples into our custom bead barcode clustering algorithm. The purpose of this step (explained in greater detail below) is to identify clusters of bead barcodes that share more UMIs than expected by chance, indicating physical co-confinement of the beads carrying these barcodes in the same droplet, and merge them, so as to avoid overcounting the same cell. The output of this algorithm is a new whitelist, this time with bead barcodes from the same droplet grouped together.
10. We repeat steps 4-7 above using the new whitelist, this time grouping together all reads that came from any bead barcode in the same droplet: umi_tools extract --error-correct-cell --bc-pattern=NNNNNNNNNN --bc-pattern2=CCCCCCCCCCCC --filter-cell-barcode -- whitelist=[NEW_WHITELIST].
11. We generate a count table of UMIs per probe per droplet: umi_tools count --per-gene -- per-contig --per-cell.

### Bead-barcode clustering and merging

We developed a computational approach to detect and resolve (1) bead barcode mutations, (2) bead multiplets, and (3) large droplets (see **Fig. S3**).

1. “Bead barcode mutations” refer to two or more bead barcode sequences that were generated from the same original bead, through bead synthesis errors or early-cycle PCR errors.
2. “Bead multiplets” represent cases where a single droplet in emulsion contains 2 or more barcoded beads, which occurs frequently by chance because we load beads into emulsions with a Poisson λ=0.4-1 (>25% of droplets with at least 1 bead will have more than 1 bead).
3. “Large droplets” occur rarely via merging during the emulsion generation process, leading to emulsified droplets that contain many (up to 100s) beads and many cells.

In each of these cases, we could spuriously overcount cells if we treat each bead barcode as representing one unique cell. In the last case, we could mistakenly combine data from many single cells into one measurement.

To detect each of these three cases, we leverage the UMIs encoded in the HyPR probes. Specifically, we identify and evaluate cases where different bead barcodes are associated with the same set of UMIs — corresponding to cases where the HyPR probes in one or more cells were amplified by primers with different bead barcodes. (This deduplication approach is not possible for whole-transcriptome scRNA-seq methods, where the UMI and bead barcode are encoded in the same primer, and is conceptually similar to a deduplication approach recently described for single-cell ATAC-seq^44^).

To implement this approach, we construct a graph in which nodes represent bead barcodes and edge weights are set equal to the fraction of overlapping UMIs between the pair of bead barcodes. We then convert these edge weights into binary 0/1 connections based on whether a pair of barcodes has more shared UMIs than expected by chance. Specifically, we keep edges between bead barcodes if the fraction of shared UMIs exceeds a given threshold, typically 0.5% in our experiments (we manually adjust this threshold as needed based on number of probes and sequencing depth). At this threshold, two bead barcodes would be merged if they each have 1000 UMIs and share at least 5; this would occur by chance <0.4% of the time, assuming approximately 1,000,000 (4^10^) possible UMIs in the HyPR-seq probes. We find clusters in this graph using a connected components algorithm (as implemented in scipy.sparse.csgraph), and return a whitelist (for use by UMI-tools) mapping each droplet to one or more bead barcodes.

In experiments with stably integrated barcodes detectable by HyPR (see below), we found that this approach increased the fraction of bead barcodes we could confidently assign to one cell. By confining our analysis to droplets with one (or only a few) beads, we were able to increase our assignment rate of cell (guide) barcodes to bead barcodes to over 80% (up from 50%), suggesting a similar improvement in data quality in experiments without ground truth cell barcodes.

### Estimating droplet occupancy rates

In order to estimate the occupancy rates of both beads and cells per droplet, we model the observed distribution (of beads or cells per droplet) as a zero-truncated Poisson distribution and fit the lambda parameter by maximum-likelihood estimation. This involves computing the sample mean (the empirical estimate of beads or cells per droplet) and solving for the occupancy rate:

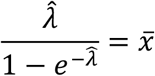

For the estimate of beads per droplet, we use the clusters from the bead-barcode deduplication procedure (described above). Specifically, we remove clusters corresponding to “large droplets” (more than 5 bead barcodes) and remove bead barcodes within clusters that are likely synthesis/PCR/sequencing errors (barcodes with Hamming distance 1 to anything else in the cluster). We use this corrected distribution of bead barcodes per droplet to compute a sample mean and plug into the equation above to solve for the occupancy rate.

For the estimate of cells per droplet, we use data from experiments where our cells were expressing one of 16 “detection barcodes”, or unique RNAs detectable with HyPR-seq probes that differ between cells. By counting the number of unique detection barcodes we see per droplet, we can infer the number of cells per droplet. (Cases where two cells with the same detection barcode occupy the same droplet will be relatively rare, approximately 1/16 x the doublet probability of ~1%.) We compute the distribution of cells per droplet by counting the number of detection barcodes in each droplet with >10 UMIs (ignoring the 10% of cells with 0 detectable barcodes), convert to a sample mean, and substitute into the equation above.

### Software

| Package | Version | Reference |
| --- | --- | --- |
| python | 3.5.1 |  |
| matplotlib | 3.0.3 | 45 |
| numpy | 1.18.0 | 46 |
| pandas | 0.24.2 | 47 |
| scikit-learn | 0.20.3 | 48 |
| scipy | 1.2.1 | 49 |
| statsmodels | 0.9.0 | 50 |
| umi-tools | 1.0.0 | 51 |
| snakemake | 5.5.3 | 52 |
| bowtie | 0.12.7 | 53 |
| samtools | 1.8 | 54 |
| FASTX-Toolkit | 0.0.13 | http://hannonlab.cshl.edu/fastx_toolkit/index.html |
| fastp | 0.20.0 | 55 |
| kallisto | 0.43.0 | 56 |
| monocle2 | 2.4.0 | 57 |
| seaborn | 0.9.0 | https://seaborn.pydata.org/index.html |
| scanpy | 1.4.1 | 58 |

### Cell Culture

#### K562

The immortalized myelogenous leukemia K562 cell line was obtained from ATCC (ATCC, CCL-243). Cells were maintained in RPMI 1640 Medium (Corning, 10-040-CM) supplemented with 10% fetal bovine serum (Life Technologies, 16140071) and 1% penicillin-streptomycin (Life Technologies, 15140163). Cell lines were cultured at 37°C with 5% CO_2_ and were subcultured twice a week by aspirating off all cell culture medium except 0.5 ml in a T-75 flask and 1 ml in a T-175 flask, and refreshing the flasks with 13 ml and 49 ml of medium respectively.

#### THP1

The human monocyte THP1 cell line was obtained from ATCC (ATCC, TIB-202). Cells were maintained in RPMI 1640 Medium (Corning, 10-040-CM) supplemented with 10% fetal bovine serum (Life Technologies, 16140071) and 1% penicillin-streptomycin (Life Technologies Inc., 15140163). Cell lines were cultured at 37°C with 5% CO_2_ and were subcultured twice a week by aspirating off all cell culture medium except 0.5 ml in a T-75 flask and 1 ml in a T-175 flask, and refreshing the flasks with 13 ml and 49 ml of medium respectively. For treatment experiments, cells were stimulated using cell culture medium with LPS at a final concentration of 0.5 μg/ml, diluted from a 5 mg/ml LPS stock (Millipore Sigma, L3024), for 20 hours.

### Hybridization Chain Reaction

We performed smFISH experiments using HCR. All HCR v3 reagents (probes, hairpins, and buffers) were purchased from Molecular Technologies. Thin sections of tissue (10μm) were mounted in 24 well glass bottom plates (VWR, 82050-898) coated with a 1:50 dilution of APTES (Sigma, 440140). Cells were spun at 350g for 15min onto 24 well glass bottom plates (VWR, 82050-898) coated with WGA (VWR, 80057-710). The following solutions were added to the tissue/cells: 10% formalin (VWR, 100503-120) for 15min, 2 washes of 1x PBS (ThermoFisher Scientific, AM9625), ice cold 70% EtOH at −20 2 hours to overnight, 3 washes 5x SSCT (ThermoFisher Scientific, 15557044, with 0.2% Tween-20), Hybridization buffer (Molecular Technologies) for 10min, probes in Hybridization buffer overnight, 4 15min washes in Wash buffer (Molecular Technologies), 3 washes 5x SSCT, Amplification buffer (Molecular Technologies) for 10min, heat denatured hairpins in Amplification buffer overnight, 3 15min washes in 5x SSCT (1:10,000 DAPI, VWR, TCA2412-5MG, in the second wash), and storage/imaging in 5x SSCT. Imaging was performed on a spinning disk confocal (Yokogawa W1 on Nikon Eclipse Ti) operating NIS-elements AR software. Image analysis and processing was performed on ImageJ Fiji. For K562 cells, StarSearch (https://rajlab.seas.upenn.edu/StarSearch/launch.html) was used to quantify HCR signal in tiff images processed to the same settings using ImageJ Fiji. For HCR quantification in kidney slices, we generated cell masks for PECs and podocytes using Fiji. Cell boundaries were determined by visual inspection. We then calculate the median background-subtracted intensity within each cell mask, where the background is taken as the median fluorescence intensity outside cellular regions.

### Determining HyPR-seq minimum specificity

We computed the relative HyPR-seq signal (counts per cell relative to non-targeting probes) for all probes tested in THP1 cells. (We decided to focus on our experiments in THP1 cells, since we targeted more lowly expressed genes than in K562 cells.) 16/18 genes tested had at least two probes with >10-fold signal compared to background, including all 15 genes expressed >1 TPM, which we estimate to be our detection threshold (**Fig. S4D**).

### qPCR

RNA extraction was performed according to manufacturer’s instructions using Qiagen RNeasy Plus Mini Kit (74136). cDNA was made according to manufacturer’s instructions using Invitrogen SuperScript III First Strand Synthesis (11752-050). 10% of undiluted cDNA was loaded into each RT-PCR according to manufacturer’s instructions using SYBR Green I Master (Roche, 04707516001). qPCR primers can be found in **Table S4**.

### Analysis of THP1 RNA-seq data

RNA-seq libraries were generated from 20 million THP1 cells (+/− stimulation with LPS) in three biological replicates using the Smart-seq2^2^ protocol in bulk. Reads were pseudo-aligned to a transcriptome index based on hg19 using kallisto^56^. TPM values were aggregated over all isoforms for the same gene. Gene expression changes were computed by dividing the counts for the tested genes in stimulated cells by the counts in unstimulated cells in three separate replicates, and statistics were performed on the three fold-change replicates.

### Mixing and single-cell purity experiment

Two K562 cell lines, each containing a specific detection barcode, were subjected to the standard HyPR-seq protocol. Probes for all 31 possible barcodes were added to the probe mixture during the hybridization step. At the droplet generation step, equal concentrations of each cell line (5,000 cells each) were mixed and loaded into the same well. The droplets containing the cell line mixture were then subjected to the standard PCR and downstream processing for library preparation.

### Construction of a gRNA vector detectable by HyPR

We constructed a vector (sgOpti-HyPR) capable of expressing both a gRNA and a “detection barcode” by modifying sgOpti (Addgene 85681) to insert a 400 bp fragment between the puromycin resistance cassette and WPRE. This fragment contains three 52 bp binding sites that, when transcribed into RNA, can be recognized by HyPR-seq probes (probe sequences in **Table S1**, barcode sequences in **Table S5**). The binding sites are separated by 50 bp of random sequence and are flanked by primer binding sites. gBlocks containing 31 unique detection barcodes were ordered from IDT, amplified using BrainBar-sgOpti-FWD and BrainBar-sgOpti-REV (**Table S4**), and added to sgOpti digested with MluI (NEB) using Gibson assembly. Sanger sequencing to confirm the barcode sequences was done using Seq916 (**Table S4**). Knockdown of GATA1 using gRNAs against its TSS and canonical enhancers (e-GATA1 and e-HDAC6) was identical compared to sgOpti alone, confirming the efficacy of the new plasmid (data not shown).

### Generation of K562 cell lines for GATA1 CRISPR experiment

gRNAs targeting regulatory elements in the GATA1 locus (GATA1 TSS, HDAC6 TSS, e-GATA1, and e-HDAC6) as well as non-targeting controls were cloned into the sgOpti-HyPR vector, as previously described for sgOpti^8^ (**Table S6**). Each guide was cloned independently into a separately barcoded version of sgOpti-HyPR, so guide-barcode pairings were known in advance. In these vectors, the gRNA and barcodes are located 1.2kb away from each other. To minimize viral reassortment, we prepared lentivirus for each of the 16 gRNAs separately, by plating 550K HEK293T cells on 6-well plates (Corning, Corning, NY), transfecting 24 hours later with 1 μg dVPR, 300 ng VSVG, and 1.2 μg transfer plasmid using XtremeGene9 (Roche Diagnostics, Indianapolis, IN), changing media 16 hours later, and harvesting viral supernatant 48 hours posttransfection. Stable cell lines expressing one gRNA-barcode pair were generated by separate lentiviral transductions in 8 μg/ml polybrene by centrifugation at 1200 x g for 45 minutes with 200,000 cells per well in 24 well plates. 24 hours after transduction cells were selected with 1 μg/ml puromycin (Gibco) for 72 hours, then maintained in 0.3 μg/ml puromycin. Separately infected cells were counted and pooled after selection with puromycin and KRAB-dCas9 (TRE-KRAB-dCas9-IRES-BFP, Addgene 85449) was induced with 1 μg/ml doxycycline (Millipore Sigma, D3072) for 24 hours before experiments.

### Analysis of GATA1 CRISPR experiment

HyPR-seq data was run through the pipeline as described above, with large droplets removed. Guides were assigned to cells by summing the UMIs for each unique combination of detection barcode probes and requiring a 6-fold excess of UMIs for the most abundant combination compared to the next most abundant (>80% success rate of assigning perturbations to cells). Cells that had silenced the CRISPRi machinery were excluded by removing the bottom 15% of cells by BFP expression (fraction of total UMIs, **Fig. S5C**). GATA1 UMIs were normalized by GAPDH UMIs in the same cell, and knockdown was computed by averaging the normalized GATA1 expression within all cells assigned to a particular guide. Differential expression testing was done with a two-sided T-test, comparing to all cells with negative control guides.

### Power calculations for HyPR-seq and 10x Genomics Chromium 3’ scRNA-seq

We computed power curves for HyPR-seq and 10x Genomics Chromium 3’ scRNA-seq using a simulation and permutation framework based on experimental data. The power for a screen depends on many factors: the effect size of the element on gene expression, the sequencing depth of a particular experiment, the number of UMIs per gene per cell, the total number of perturbed elements, the number of guides per cell, and others. Here we computed power based on two summary variables: effect size and number of perturbed cells.

We computed power to detect 10%, 25% and 50% effects on each gene as follows. For each combination of effect size and “number of cells measured per perturbation” (N), we:

1. Simulated a dataset in which N cells are randomly selected to be ‘perturbed’ and M - N cells are not perturbed (M = total cells from a given single-cell dataset, M = 200,000 for 10x Genomics Chromium 3’ scRNA-seq and 5,000 for HyPR-seq). This is to simulate an experiment with an excess of non-targeting negative control gRNAs.
2. For each non-perturbed cell, we estimate its UMI counts by sampling from a Negative Binomial distribution for each gene. The mean and dispersion for each NB are fit from experimental data (UMIs per cell for that gene across all cells in the given dataset).
3. For each perturbed cell, the mean for the NB comes from a 10/25/50% decrease from the non-perturbed mean, and the dispersion is the estimated dispersion at the decreased mean as expected from the mean-dispersion fit across all genes in the experimental data.
4. Given this simulated dataset, we test for differential expression in perturbed cells using Monocle2^57^. We select differentially expressed genes at an FDR of 5% across all tested genes in the dataset.
5. We repeat the above steps 100 times, and compute power as the number of significant tests (per gene) divided by 100.

We ran these simulations for 37 genes measured in a HyPR-seq dataset from K562 cells, spanning a range of expression levels (1.5-368 UMIs/cell in HyPR-seq, corresponding to >0.7 TPM in RNA-seq and >0.01 UMIs/cell from scRNA-seq). We plotted power averaged over all 37 genes at each (effect size, cells measured) tuple.

### Kidney Single Cell Dissociation

All animal work was done according to IACUC protocol #0061-07-15-1. Two BTBR *wt/wt* and two homozygous BTBR *ob/ob* male mice at 12 weeks of age (Jackson Laboratories, 004824) were anesthetized using 4% Isoflurane (029405, Henry Schein Animal Health), then transferred to a nose cone supply for the duration of the procedure. Performing a combination of blunt dissection and scissor-assisted dissection techniques, the visceral organs were exposed before opening the thoracic cavity to show the thoracic organs. The right atrial-chamber was lacerated with scissors before inserting a 27-gauge scalp vein butterfly needle (Excel International, 14-840-38) into the left ventricular chamber and perfusing with 10-20 mls of ice-cold 1X PBS (ThermoFisher Scientific, 10010023) using a variable-speed peristaltic pump (VWR, 70730-062) until the heart stops beating and the liver blanches. The kidneys are removed, cut in half, and placed in ice-cold 1X PBS before the renal capsule is removed and the tissue is stored on ice in 1X PBS. Before, prepared 2.5 mg/ml Liberase TH by diluting 10 mg of Liberase TH (Sigma Aldrich, 5401135001) in 5 ml of DMEM/F12 (Life Technologies, Inc., 11320033) and stored at −20°C. Right before manual dissociation, thawed and diluted one 100 ul aliquot of 2.5 mg/ml Liberase TH with 0.9 ml of DMEM/F12 per half kidney to make 1X Liberase TH in DMEM/F12. Each half kidney was transferred to a petri dish and manually dissociated using tweezers and a razor blade, then resuspended well using a 1 ml pipette tip in 1 ml of 1xLiberase TH in an Eppendorf tube. Tubes were incubated in a thermomixer at 600 rpm at 37°C for 2h; and were gently and thoroughly pipetted up and down with a 1ml pipette tip every 10 min. Two kidney halves from the same mouse were combined and homogenized using 40 passes of a dounce homogenizer on ice. This was repeated for the other kidney before combining all samples from the same mouse in a 50 ml conical and adding 40 ml 10% FBS RPMI media to stop the digestion. The media was prepared prior by adding 50 ml FBS (Life Technologies Inc., 16140071) to a 500 ml bottle RPMI (Corning, 10-040-CV) media. Centrifuged at 500xg for 5 min at room temperature (RT). Aspirated off the supernatant and resuspended pellet in 4 ml of Red Blood Cell Lysing Buffer Hybri-Max (Sigma-Aldrich, R7757-100ML). Centrifuged at 500xg for 5 min at RT. Aspirated off the supernatant and resuspended pellet in 1 ml Accumax (Stemcell Technologies, 07921) for 3 min at 37°C. Added 20 ml of 10% FBS RPMI to neutralize the Accumax and centrifuged at 500xg for 5 min at RT. Aspirated off the supernatant and resuspended the pellet in 4 ml 0.4% BSA/PBS, prepared by dissolving 80 mg of Bovine Serum Albumin (Sigma-Aldrich, A9418-10G) in 20 ml of PBS the day before. Filtered solution through a 30 μm filter (Corning, 351059), then through a 20 μm pluristrainer (Pluriselect, 43-50020-01). Diluted the sample to assess the cell number and viability using trypan blue (Sigma-Aldrich, T8154-100ML) and a cellometer (Nexcelom Bioscience, Cellometer Auto T4). Samples were kept on ice before proceeding to fixation in the HyPR-seq protocol.

### Computational analysis of kidney HyPR-seq data

HyPR probes for kidney cells were chosen based on canonical marker genes for PCTs, DCT/TAL, podocytes, mesangial, endothelial, macrophages, CD-IC, and CD-PC cell types.

#### Filtering and normalization

For initial quality control of the data matrix consisting of 37,725 (18,853 WT + 18,872 DKD) cells and 32 genes, we filtered out cells that were more than two standard deviations from the mean in each of total UMIs per cell (1,527 cells), expressed genes per cell (1,325 cells), and expression level of a housekeeping gene (*B2m*, 2,017 cells). We also filtered out the cells that had non-zero expression of “GFP” and “GAPDH_human” probes, since these genes should not be expressed in mouse cells (439 cells). 14.1% of the cells (5,308 cells) were filtered out in these steps. We did not perform batch correction, since the difference of the distribution of total count and the number of expressed genes per cell was minimal between the batches in the same biological condition (Kolmogorov-Smirnov test D statistics <0.1, **Fig. S7A**). For the remaining 32,417 cells, we scaled the total counts per cell to 10,000 to enable comparisons between cells with different total counts. Next, we converted the counts into log scale (log2+1) and scaled the expression for 32 genes to be unit (mean=0, var=1) in order to avoid overweighting highly expressed genes in the clustering step. This scaled expression level was used for defining the cell identities, as well as for the heatmaps and dot plot.

#### Clustering

We projected the data into 6-dimension principle component space, and performed Leiden’s clustering method with 100 nearest neighbors and resolution=0.7 to identify distinct cell populations. Next, in order to put the clusters into biological context, we first manually combined sets of clusters whose difference in the expression pattern cannot be characterized by distinct genes (and therefore thought to be driven by technical artifacts). Specifically, six distinct clusters with >50% non-zero expression in each of seven genes that are commonly expressed in PCT (*Aqp1, Atp11a, Enpp2, Lrp2, Miox*, Napsa and *Slc34a1*) and <20% non-zero expression of any other marker genes, were combined and defined as “PCT” (We assumed further characterization of PCT subtype within each of these clusters is not possible without additional probes). Two distinct clusters with >50% non-zero expression of Wt1 and <20% non-zero expression of *Nphs2, Synpo, Ptn, Itga8* and *Ctgf* were combined and defined as “PEC”. Two clusters with >80% non-zero expression of each of three endothelial cell marker genes (*Ehd3, Kdr* and *Pecam1*) were combined and defined as “Endo”. We also manually removed two clusters thought to be cell duplets, characterized by the expression of combinations of marker genes of different distinct cell types that were not validated in HCR staining (*e.g.*, we removed clusters with >80% non-zero expression of both PCT marker genes described above and one of mesangial cell marker genes: *Ptn* and *Itga8*). The post-clustering filtering steps described above reduced the number of cells from 32,417 to 29,125, and the number of clusters from 21 to 11. We do not fully exclude the possibility of merging or removing rare novel cell populations in these steps. The full cluster identity before the filtering steps is available in **Fig. S7**.

#### Visualization

Low dimension embedding using UMAP was performed with the default parameter specified in scanpy package, with seed=1. Six genes (housekeeping genes: *Gapdh, B2m, Rpl13a, Hprt*, as well as 2 negative control probesets: *GAPDH* (human) and GFP) with the lowest variance of the expression were removed in the visualization. For the quantification of the correlation with scRNA-seq clusters, we calculated the mean expression of each gene in each cluster (both for HyPR and scRNA-seq), and calculated the Pearson correlation. Seven genes (*Atp11a, Atp6vo1b2, Enpp2, Kdr, Muc1, Pecam1, Slc22a7*) were excluded because they were missing in scRNA-seq data, presumably due to their low expression.

#### Cell type frequency plot

The fold change of cell types between wild-type (WT) and diabetic (DKD) mice was calculated from the relative fraction of all cell types within each replicate, averaged for the wild-type and diabetic mice. The error was calculated by:

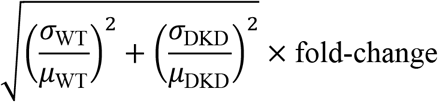

## Acknowledgements

This work was supported by the Broad Institute, an NIH Early Independence Award (1DP5OD024583 to F.C.), the Schmidt Fellows Program at the Broad Institute (F.C.), and an NIH Pathway to Independence Award (1K99HG009917 to J.M.E.). J.M.E. was supported by the Harvard Society of Fellows. S.G.R. was supported by the Hertz Graduate Fellowship and the National Science Foundation Graduate Research Fellowship Program (award 1122374). Q.W. was supported by the Nakajima Foundation Scholarship. The authors thank Jason Buenrostro, Christoph Muus, Aviv Regev, Arjun Raj, Vijay Sankaran, and Tung Nguyen for discussions.

## Competing interests

F.C., J.M.E., J.L.M., V.S., and S.R. are inventors on patent applications filed by the Broad Institute related to this work (62/676,069 and 62/780,889). E.S.L. serves on the Board of Directors for Codiak BioSciences and Neon Therapeutics, and serves on the Scientific Advisory Board of F-Prime Capital Partners and Third Rock Ventures; he is also affiliated with several non-profit organizations including serving on the Board of Directors of the Innocence Project, Count Me In, and Biden Cancer Initiative, and the Board of Trustees for the Parker Institute for Cancer Immunotherapy. He has served and continues to serve on various federal advisory committees.

## SUPPLEMENTAL NOTES AND FIGURES

### Note S1. Design considerations for HyPR-seq for sensitive, multiplexed detection of chosen RNAs in tens of thousands of single cells

We set out to develop a method to quantify selected RNAs in single cells. To accomplish this, we designed a method that could (i) detect chosen transcripts with good per-molecule sensitivity, (ii) allow multiplexed detection of up to 100 transcripts, (iii) allow detection of non-polyadenylated transcripts or RNA splice isoforms, (iv) measure up to 100,000 cells with relative ease, and (v) achieve these capabilities with costs on the order of a few cents per cell. In the following sections, we discuss how we designed HyPR-seq to achieve these goals. We also discuss conceptual differences between HyPR-seq and existing methods for targeted single-cell quantification of RNA expression.

#### HyPR-seq achieves high detection rates for individual transcripts of interest with minimal reads per cell

A key goal of our approach is to obtain sensitive quantification of a given RNA of interest in order to perform differential expression tests, detect low-abundance marker genes in single cells, and/or examine gene expression dynamics. To accomplish this goal, we must consider two related parameters: the absolute detection rate (*i.e.*, the fraction of RNA molecules in a given cell that would be detected given infinite sequencing depth) and the total number of sequencing reads required to achieve a given number of counts per cell.

We designed HyPR-seq to address both of these considerations. Toward achieving a high detection rate, we noted that single-molecule fluorescence *in situ* hybridization (smFISH) can achieve very high (approaching 100%) sensitivity for detecting individual RNA transcripts in cells^1–5^. The high sensitivity of smFISH likely results from the fact that (i) each RNA is targeted with many individual single-stranded DNA probes, providing several opportunities to visualize each molecule, and (ii) that smFISH involves nucleic acid hybridization for detection, which is highly efficient and requires minimal enzymatic or processing steps that could lead to RNA loss. While smFISH provides highly efficient readouts of particular RNAs, it does not provide all of the capabilities needed for our goals. Initial smFISH methods could only examine a handful of RNAs at a time by imaging fluorescent probes in different colors^1–5^. Recent approaches — such as MERFISH, seqFISH, and *in situ* RNA sequencing — have increased multiplexing using combinatorial barcoding or direct sequencing of hybridized probes, but they typically require specialized equipment and long imaging times^6–8^.

We developed HyPR-seq to combine the high detection rate of smFISH with the throughput and ease of next-generation DNA sequencing. To do so, we adapted the probes designed for hybridization chain reaction (HCR) version 3 for a sequencing-based readout^5,9^. Similar to HCR, we design pairs of DNA probes against specific sequences along one or many RNAs of interest. Only paired probes that specifically bound adjacent to each other on the target transcript allow capture of the B1H1 hairpin oligo (**Fig. S1**), ensuring high specificity. The B1H1 oligo in turn serves as a template for the “readout” oligo which is ligated to the 5’ end of the UMI containing target probe, resulting in a PCR amplifiable target probe template. On a per-probe basis, we observe an estimated 20% sensitivity for detecting individual RNA molecules, and by designing multiple probe pairs against a target transcript, as in smFISH, we can achieve total sequencing counts per cell similar to the number of molecules in a cell.

This targeted approach also enables us to reduce the number of sequencing reads required to achieve a given number of counts per cell. Lowly expressed transcripts might be present at a level of ~1-10 transcripts per million (TPM), indicating that whole-transcriptome single-cell RNA sequencing experiments must sequence at least millions of reads in order to ensure robust detection in a given cell or cell population. This cost of sequencing often makes it impractical to use these approaches to study individual transcripts of interest beyond those that are highly abundant in a cell. To address this issue, HyPR-seq allows the user to sequence only a subset of chosen transcripts, thereby reducing the sequencing cost required to observe a given number of counts per transcript per cell by several orders of magnitude. In a given experiment, only 100-10,000 reads per cell might be required to detect such a lowly expressed transcript, depending on the total number and expression levels of the other genes selected for the experiment.

Several recent approaches provide similar ability to focus sequencing power on individual transcripts of interest, for example by using hybrid selection or multiplexed PCR to enrich selected transcripts from whole-transcriptome 10x scRNA-seq libraries^10^. HyPR-seq extends the capabilities of these methods in several ways: HyPR-seq sequences only the selected transcripts (there is no background from other RNAs in the cell, because only the properly hybridized and ligated probes provide a template for PCR), enables tuning the numbers of reads on a per-transcript basis by including more or fewer probes for a given transcript, and introduces capabilities for multiplexed detection non-polyadenylated RNAs (see below).

#### HyPR-seq allows multiplexed detection of up to 100s of transcripts

A key design goal for HyPR-seq is to allow multiplexed detection of up to 100s of RNA transcripts, enabling flexible research designs for applications including CRISPR screens (where one might want to study the effects of an enhancer on several nearby genes) and studies of primary tissues (where one might want to use 100s of marker genes to quantify the frequencies of dozens of cell types in a tissue). In HyPR-seq, the user can achieve this multiplexing by simply adding different sets of probes to the experiment, all of which have the same universal PCR handles and are quantified in parallel through high-throughput sequencing. In this study, we perform experiments involving 100-250 probes in a single experiment (200-4,000 UMIs per cell), indicating that HyPR-seq could detect up to hundreds of genes in parallel. Experiments could be scaled up to even larger numbers of probes.

#### HyPR-seq enables sensitive detection of non-polyadenylated transcripts

We designed HyPR-seq to allow detection of arbitrary combinations of both polyadenylated and non-polyadenylated transcripts, thereby enabling single-cell studies to detect specific splice isoforms and to compare the dynamics of gene intron and exon abundance, among other applications. HyPR-seq achieves this by detecting RNA using DNA probes, which can be designed to hybridize to arbitrary 52 bp RNA sequences without relying on 5’ or 3’ capture. In this study, we demonstrated that HyPR-Seq can detect both exonic and low abundance intronic RNA species, enabling precise and temporal quantitative measurements of transcription rates.

Existing single-cell RNA-seq approaches detect some signal from introns with endogenously encoded polyadenylation tracts using standard oligo-dT primers^11^ and can extract signal from other non-polyadenylated transcripts via the use of target-specific or semi-random reverse transcription primers^12–14^. However, these approaches are limited by cell throughput or by multiplexing ability. To our knowledge, HyPR-seq provides the first approach to enable sensitive and multiplexed detection of chosen non-polyadenylated transcripts in thousands of single cells.

#### HyPR-seq can profile up to 100,000 single cells with relative ease and low cost

We designed HyPR-seq to enable profiling up to 100,000 single cells in a single experiment. Such capabilities are essential for applications involving detecting rare cell types in a tissue, performing large-scale perturbation screens, and detecting subtle changes in the expression of individual genes. To achieve this high cell throughput, we leverage microfluidic droplet technology, which has emerged as a powerful tool for single cell analyses due to their throughput, single and fast workflow and consistent data quality. In particular, HyPR-seq uses a commercially available microfluidics instrument — the Bio-Rad QX200 Digital Droplet PCR (ddPCR) system — to generate nanoliter-scale droplets that are stable and compatible with subsequent PCR. In this system, one lane (similarly sized to one well of a standard 96-well plate) generates ~20,000 droplets, into which we Poisson-load cells and barcoded beads to obtain 500-2,000 cells per well. The Bio-Rad system generates droplets for 8 lanes at a time in the span of minutes, allowing serial loading of hundreds of wells to profile tens to hundreds of thousands of cells, either from a single experiment or distributed amongst many individual experiments.

#### HyPR-seq enables low cost single cell transcript measurements

We designed HyPR-seq to enable measuring hundreds of thousands of single cells in a cost-effective manner. Experimental costs for whole-transcriptome scRNA-seq approaches (such as 10x Genomics Chromium scRNA-seq) involves two major categories: (i) the reagent costs required to generate the libraries and (ii) the sequencing costs needed to read out the entire transcriptome. HyPR-seq reduces both of these costs, by (i) using off-the-shelf reagents (Bio-Rad ddPCR machine, ssDNA oligos from IDT, barcoded beads from Chemgenes, etc.) instead of proprietary reagents for emulsion generation and library preparation and (ii) focusing sequencing power on the small number of transcripts of interest. For the experiments in this study, we estimate that the total cost was less than 5 cents per cell, including probe, reagent, and sequencing costs. Furthermore, due to the increased transcript counts per cell for selected RNAs, HyPR-seq also reduces the number of cells required for certain applications, such as well-powered measurements of the effects of CRISPR perturbations (see main text). As such, for certain experiments such as large-scale enhancer screens, HyPR-seq can reduce the total cost by over 100-fold versus whole-transcriptome scRNA-seq.

In summary, HyPR-seq provides unique capabilities for targeted, high-throughput, sensitive, and cost-effective RNA detection in single cells. We anticipate this approach will enable new types of experiments, including to quantify the dynamics of RNA expression and splicing, map the functions of *cis-*regulatory elements, and perform high-throughput screens *in vivo* to understand the effects of perturbations on tissue composition.

**Fig. S1.**
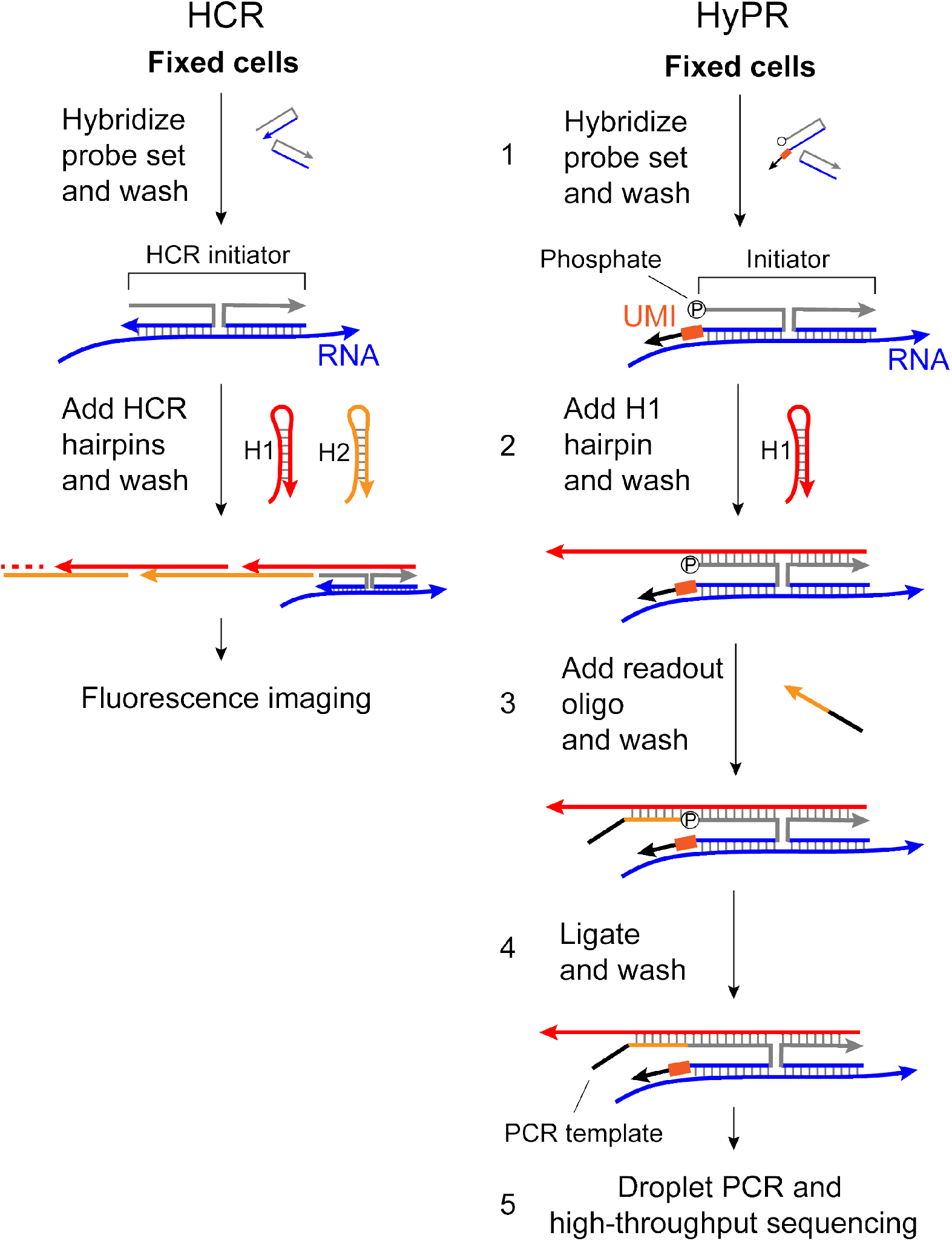
Adapting HCR probes for HyPR-seq. In both protocols, the first step (1) involves hybridizing a pair of probes that recognize a specific target site on RNA, which anneal adjacent to one other and create an initiator sequence. In the second step (2), HCR involves adding two metastable hairpins (H1 and H2) that anneal to each other to form a long chain containing fluorophores that are then detected by fluorescence imaging. HyPR-seq modifies this procedure in several ways. (1) One of the initiator probes contains a UMI (dark orange), a 3’ PCR primer binding site (black), and 5’ phosphate. (2) Only one hairpin (H1) is added. (3) A readout oligo (a shortened version of the H2 hairpin with a 5’ PCR primer binding site) is added and then (4) ligated to the left initiator probe. The resulting ligated ssDNA product is then amplified during a PCR step in droplets to generate libraries for high-throughput sequencing.

**Figure S2.**
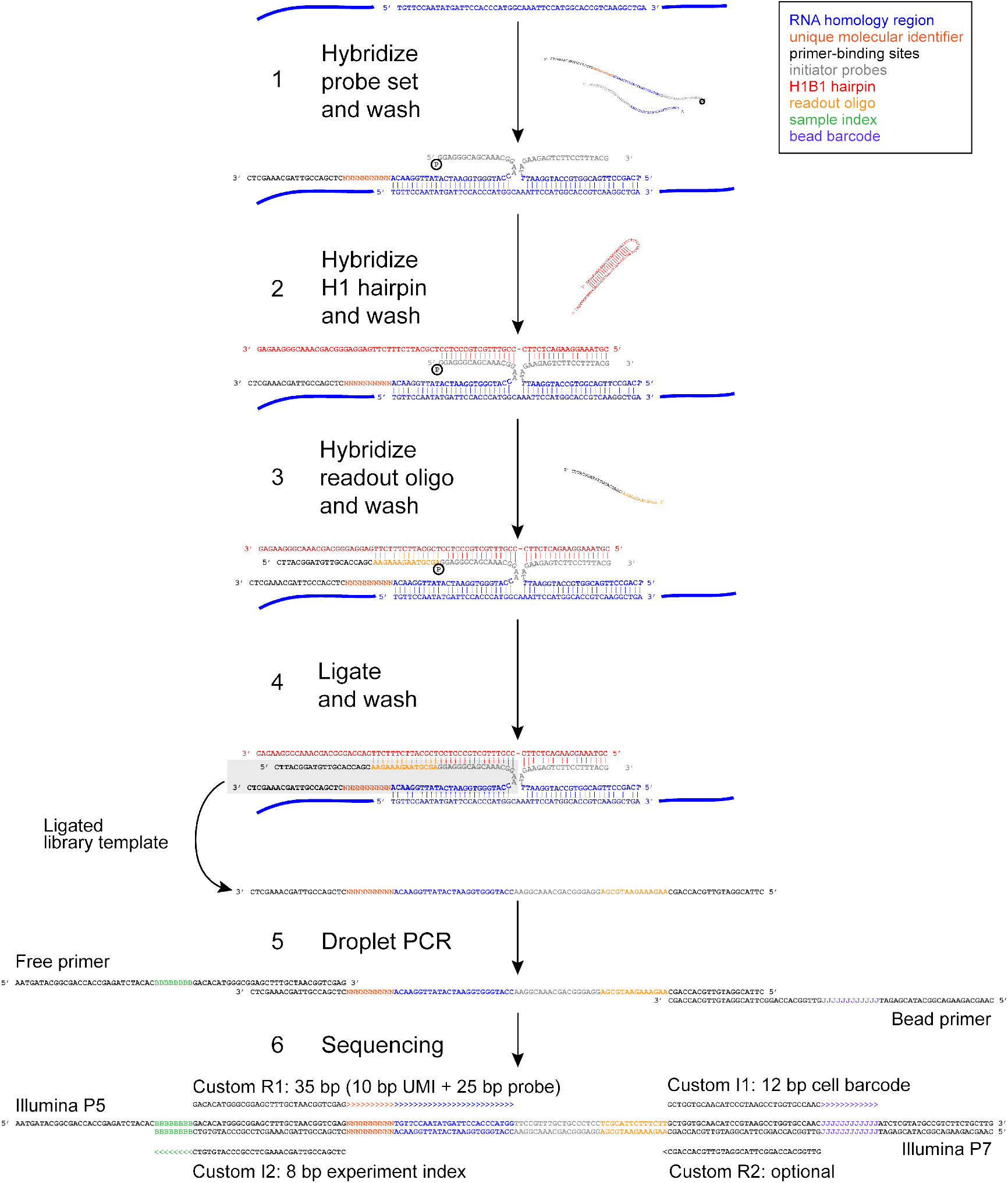
HyPR-seq probes and sequencing amplicons. (1) We hybridize HyPR-seq initiator probes to mRNA transcripts in the cell (these sequences shown here represent GAPDH probe 7, **Table S1**). The left initiator probe includes a 5’ phosphate group, UMI, and 3’ primer binding site. The two initiator probes bind to the mRNA with a 2 bp gap between 5’ and 3’ probe binding sites. (2) Following extensive washes, we hybridize H1 to the initiator probes. The presence of the paired initiator probes allows H1 to unfurl from its metastable hairpin structure and hybridize to the initiators. (3) After washing out excess H1, we add the readout oligo, which hybridizes to H1. (4) After washing, we ligate the readout oligo to the left initiator probe to form the library template. (5) We emulsify cells in droplets with DNA-barcoded beads and free primer in solution containing the sample index. The final dsDNA product shows the final sequencing amplicon with primers for the Illumina NextSeq 500 platform.

**Figure S3.**
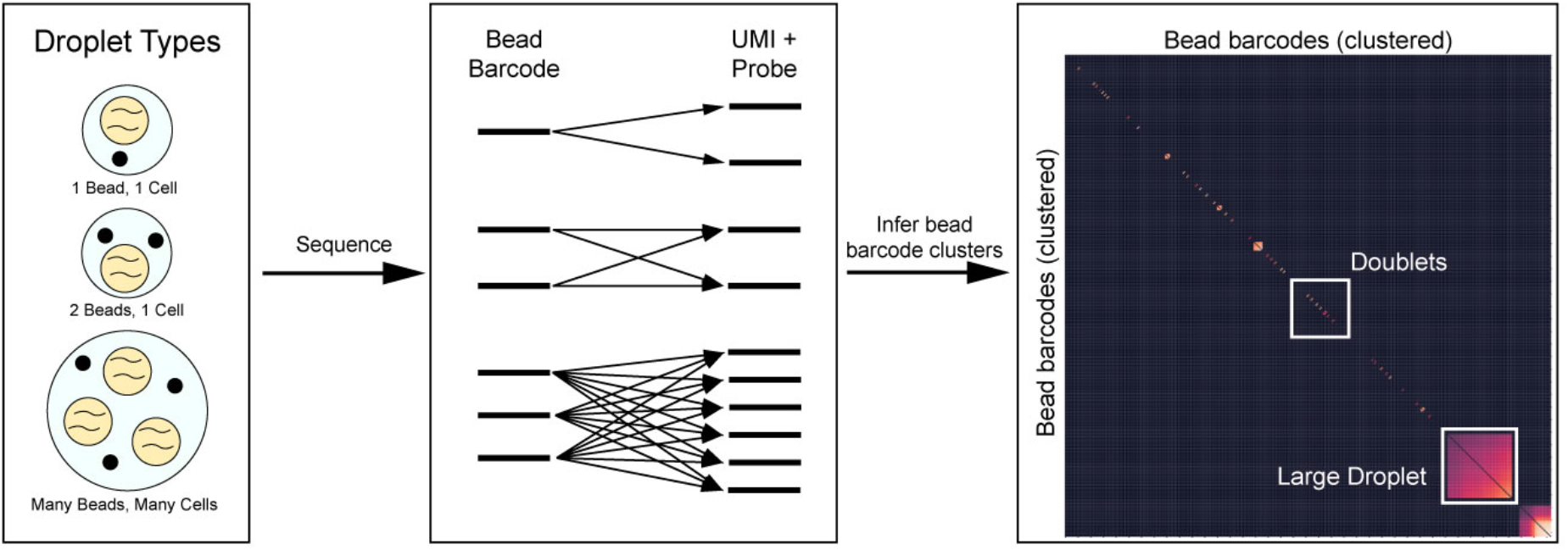
Bead barcode deduplication. Schematic of the computational procedure for identifying bead barcodes that come from the same droplet. Because we Poisson load beads into droplets, there can be cases when one droplet ends up with more than one bead (left). In such cases, primers from multiple beads will amplify HyPR-seq probes sequences with the same UMIs (center). This allows us to cluster bead barcodes by shared UMIs and identify bead doublets (two beads with one cell) and some large droplets (multiple beads and cells, right). In the cluster diagram at right, each pixel shows the fraction of overlapping UMIs for a pair of bead barcodes (entries on the main diagonal are ignored). Pairs of squares immediately off the diagonal indicate the presence of bead doublets (two beads that share many UMIs with each other), while bigger squares result from “large droplets” containing many beads and many cells.

**Figure S4.**
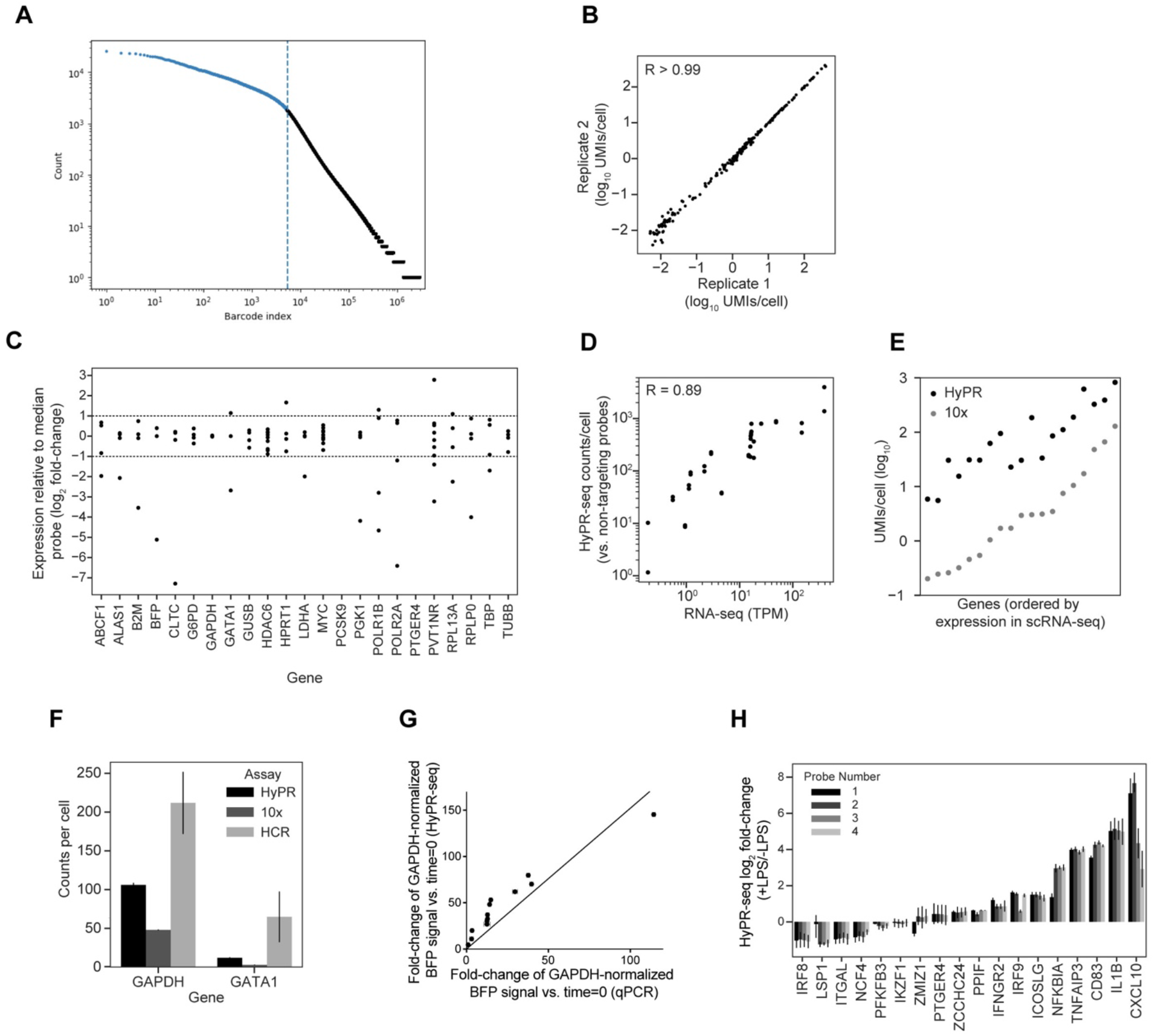
HyPR-seq QC, probe sensitivity, and quantitation. **A.** UMIs per bead barcode for the 22 gene HyPR-seq experiment in K562 cells. UMIs per cell (Y axis) are plotted in descending rank for all bead barcodes (X axis). Blue points to the left of the dotted line represent bead barcodes associated with “real” cells selected for further analysis (see Methods). **B.** Correlation between the counts per cell for each probe tested in K562 cells for two biological replicates. **C.** The expression levels of all probes tested in K562 cells are shown relative to the median expression of a probe targeting each gene. Dotted lines indicate +/− 2-fold change in expression from the median per gene. **D.** HyPR-seq signal (counts per probe relative to non-targeting probes) for 18 genes in THP1 cells (two probes shown per gene) plotted against their expression by bulk RNA-seq. 16/18 genes (and all genes >1 TPM) have at least 2 probes with >10-fold signal compared to background, **E.** UMIs per cell for 19 genes in both HyPR-seq and 10x Genomics Chromium 3’ scRNA-seq are plotted in increasing order of expression in the scRNA-seq dataset. **F.** Counts per cell for GATA1 and GAPDH in K562 cells as assayed by HCR (smFISH), HyPR-seq, and 10x Genomics Chromium 3’ scRNA-seq. **G.** Fold-change in blue fluorescent protein (BFP) RNA expression at each of 14 time points versus time = 0, as measured by qPCR (X axis) or HyPR-seq (Y axis). For both qPCR and HyPR-seq, the signal for BFP was normalized to the signal for GAPDH. Error bars represent the standard deviation of three replicates at each timepoint. **H.** Fold-change in gene expression in THP1 cells treated with LPS vs. untreated cells for each of four probes targeting 18 genes. Error bars represent the 95% CI from four biological replicates.

**Figure S5.**
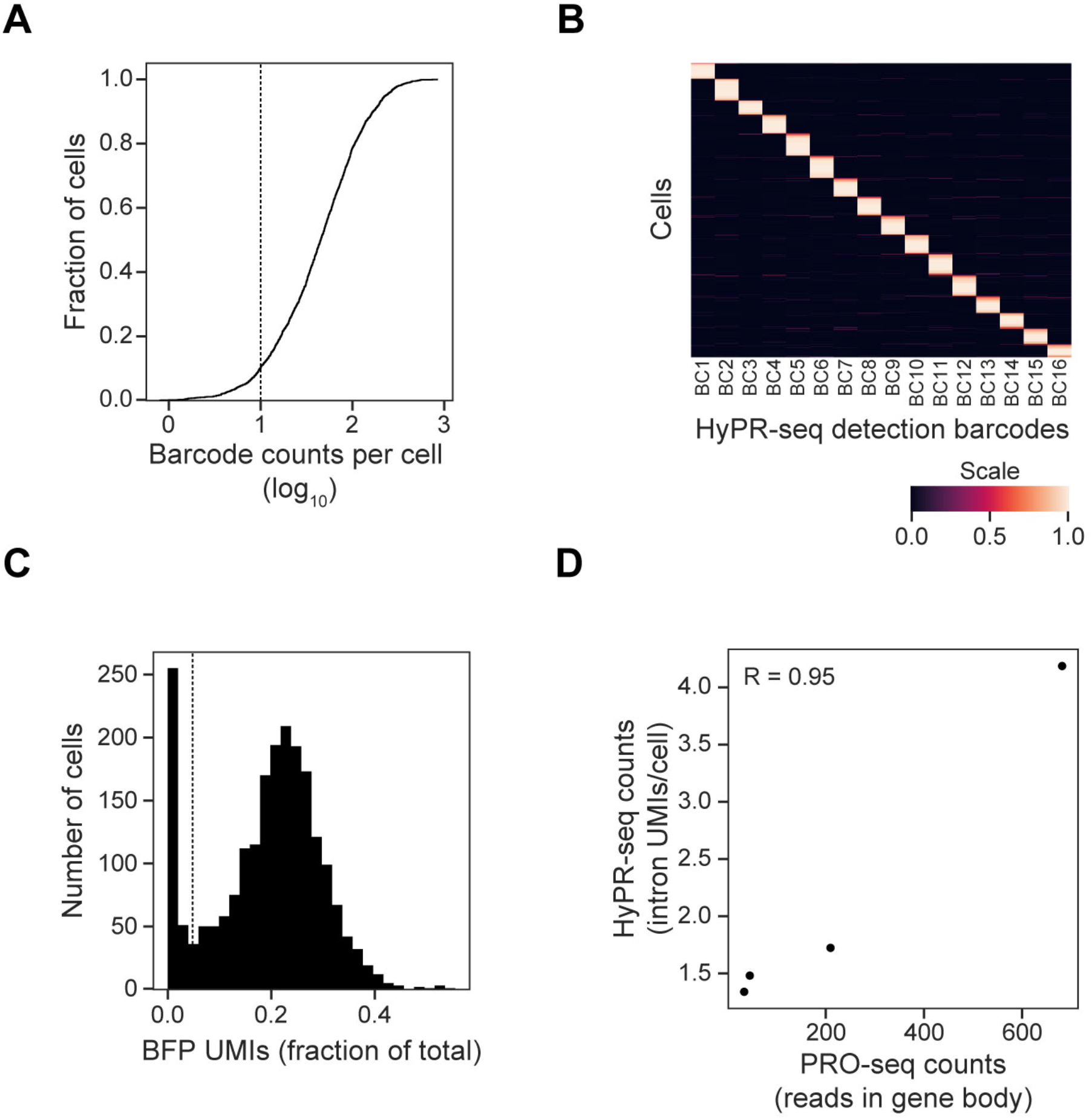
HyPR-seq detects guideRNA barcodes and non-polyadenylated gene introns. **A.** CDF of counts per cell for the HyPR-seq detection barcodes. More than 90% of cells include at least 10 UMIs for detection barcodes (dotted line). **B.** Clustered heatmap showing the HyPR-seq UMI counts of 16 detection barcodes (columns) per cell (rows) in K562 cells. Barcode counts per cell were normalized to sum to 1. **C.** Histogram of BFP counts (as a fraction of total UMIs per cell) for K562 cells expressing KRAB-dCas9-IRES-BFP in the GATA1 locus CRISPR screen. Detection of BFP indicates successful expression of the KRAB-dCas9 construct. Cells without robust KRAB-dCas9 expression were removed prior to downstream analyses by filtering out cells with under 15^th^ percentile (dotted line) expression of BFP. **D.** Correlation between transcription rates as measured by PRO-seq (X axis) or HyPR-seq using intron probes (Y axis) for four genes in K562 cells. PRO-seq signal is computed by counting reads in the gene body, and HyPR-seq counts only include the most abundant probe targeting each gene’s first intron.

**Figure S6.**
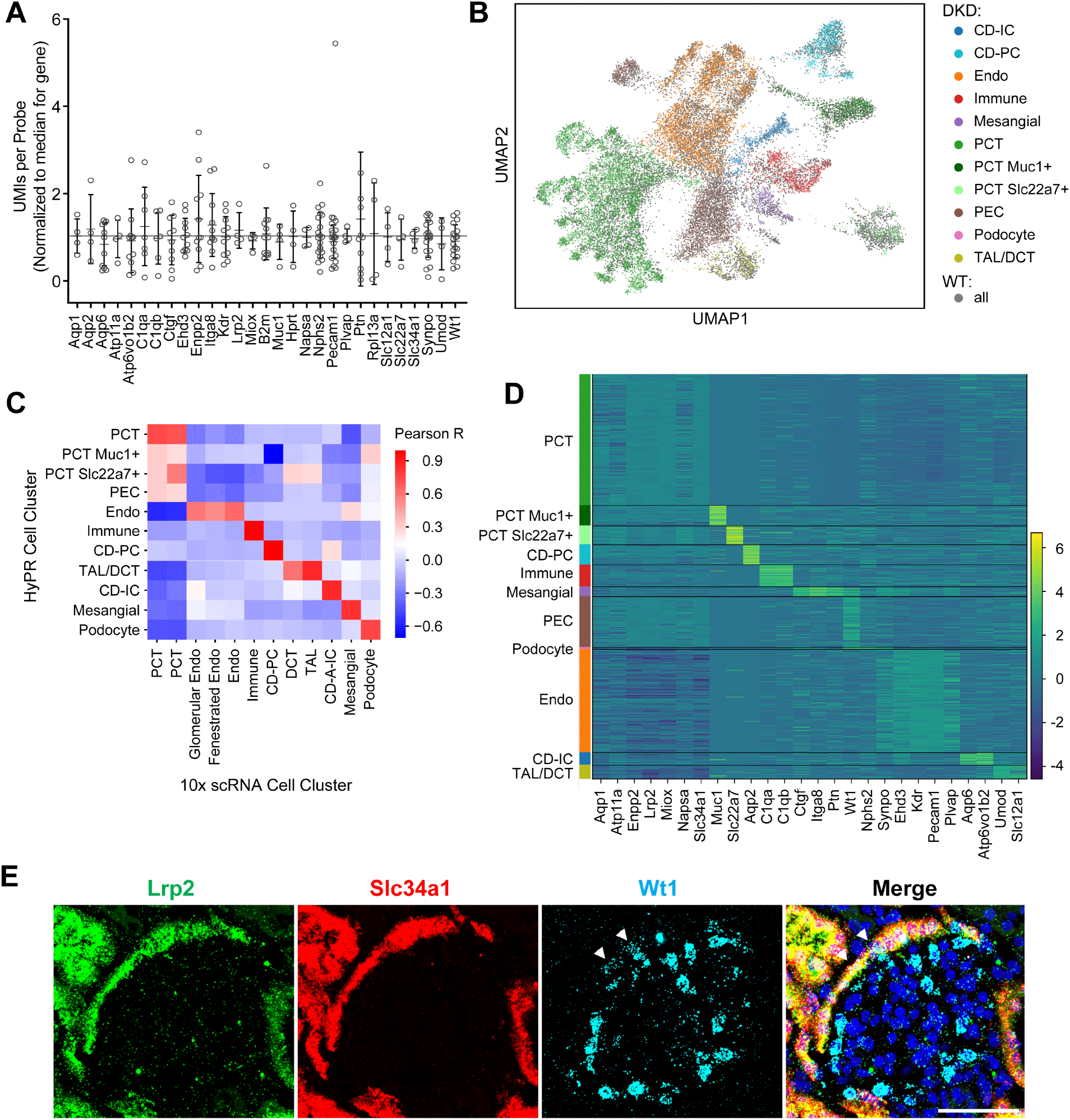
HyPR-seq detects cell types in primary murine diabetic kidney disease. **A.** Relative efficiencies of the 4-20 probes targeting each gene. Counts for each probe are normalized to the median of probes for that gene. Most probes show efficiencies within a 2-fold range of the median. **B.** UMAP visualization of 14,288 BTBR *wt/wt* cells (grey) and 14,837 BTBR *ob/ob* cells (colored) displaying 11 distinct cell types. **C.** Correlation of gene expression in HyPR-seq (rows) versus the corresponding cell populations in single-cell RNA-seq (columns). Gene expression values were averaged among cells in each population. **D.** Clustered heatmap showing the expression levels of marker genes. Cells are ordered based on the cluster identity (row). Color scale shows normalized gene expression levels (see Methods). **E.** We used HCR to visualize the localization of parietal epithelial cells (PECs), a rare cell observed in HyPR-seq data. PECs (white arrows), marked by the combination of *Wt1, Lrp2*, and *Slc34a1*, reside adjacent to the glomerulus lining the Bowman’s capsule as expected^15,16^. Scale bar: 50μm.

**Figure S7.**
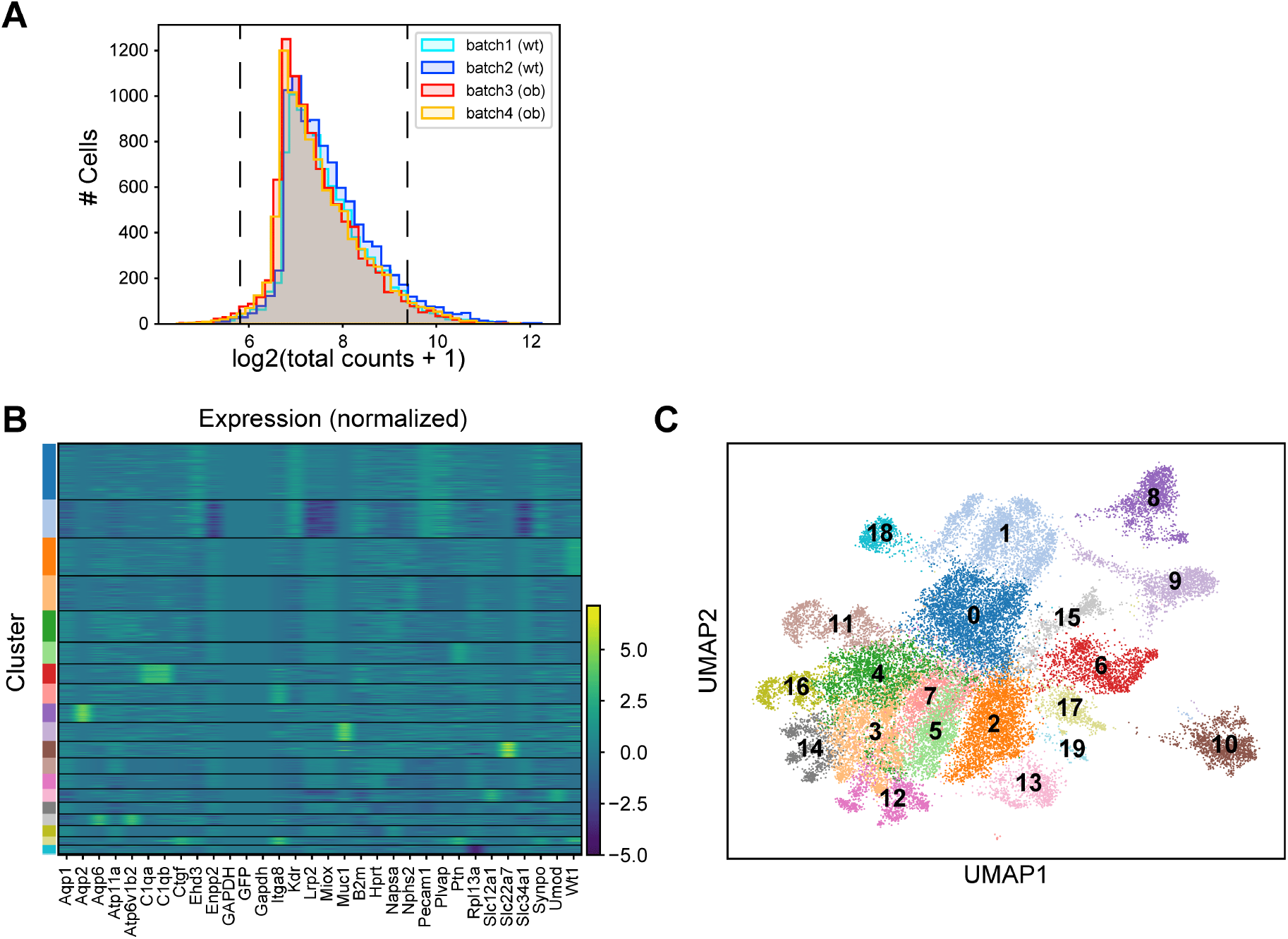
Quality control for the expression matrix. **A.** Histogram of the total expression counts, for four different batches. Black line corresponds to the +-2 standard deviation cut-off. The D statistics of the two-sample KS test between batches of the same biological replicate (*wt/wt* or *ob/ob*) were lower than 0.1, and that between *wt/wt* and *ob/ob* were lower than 0.2. **B.** Clustered heat map showing the expression level of marker genes for each of the clustered cells (row), before the removal and merging of subsets of clusters. Genes are ordered alphabetically. **C**. UMAP visualization of the cells, before the removal and merging of subsets of clusters. In (B) and (C), 32417 cells are included. From here, clusters 0 and 1 are combined as “Endo”, clusters 2 and 18 were combined as “PEC”, clusters 3, 4, 11, 12, 14, 16 were combined as “PCT”, and clusters 5 and 7 are removed to construct the final set of clusters, as described in the methods. Note that the cluster colors do not necessarily correspond to the colors in **Fig. 3** or **Fig. S6B**.

